# Antidepressants interact with an LPA_3_ receptor binding site: functional and docking studies

**DOI:** 10.1101/2025.03.18.644008

**Authors:** K. Helivier Solís, Ana I. Jardón-Ibañez, M. Teresa Romero-Ávila, Ruth Rincón-Heredia, José Correa-Basurto, J. Adolfo García-Sáinz

**Affiliations:** Departamento de Biología Celular y Desarrollo; Unidad de Imagenología, Instituto de Fisiología Celular, Universidad Nacional Autónoma de México. Ciudad Universitaria. Ap. Postal 70-600, Ciudad de México 04510. México; Laboratorio de Diseño y Desarrollo de Nuevos Fármacos e Innovación Biotecnológica, Escuela Superior de Medicina, Instituto Politécnico Nacional. Plan de San Luis y Díaz Mirón S/N, Casco de Santo Tomas, Miguel Hidalgo, Ciudad de México11340, México

**Author notes:** Correspondence should be addressed to: J. A. García-Sáinz Inst. Fisiología Celular, UNAM Ciudad Universitaria, Ap. Postal 70-600 Ciudad de México 04510 México.

**Keywords:** LPA_3_ receptors, lysophosphatidic acid receptors, antidepressants, paroxetine, imipramine, amitriptyline

## Abstract

The action of the antidepressants imipramine, amitriptyline, and paroxetine on LPA_3_ receptors was studied *in cellulo*, using receptor-transfected HEK 293 Flp-In TREx cells, and *in silico*, through docking simulations. These drugs showed a low affinity for LPA_3_ receptors with lesser efficacy than LPA (paroxetine ≈ 60% and imipramine and amitriptyline ≈ 30%). When LPA-treated cells (with the agonist present) were challenged with the antidepressants, paroxetine triggered a robust increase in intracellular calcium, whereas imipramine and amitriptyline decreased the calcium concentration below baseline values. For ERK 1/2 phosphorylation, imipramine induced a rapid and potent increase, whereas amitriptyline and paroxetine reduced ERK 1/2 phosphorylation below baseline. Similarly, imipramine produced rapid and robust ERK phosphorylation in LPA-stimulated cells, but amitriptyline decreased ERK 1/2 phosphorylation. Activation with the antidepressants leads to LPA_3_ internalization; dramatic morphological changes accompany these actions. Docking simulations showed these drugs interact with an LPA_3_ receptor pocket, denominated Upper Cavity. Although the agonist binding cavity was the same, the amino acids interacting with the various ligands were distinct due to their different chemical structure. The manuscript advances knowledge on the mechanisms of antidepressant effects on LPA_3_ receptors, which might have potential therapeutic implications.

## 1. Introduction

LPA is a lipid metabolite that mediates a large variety of functions. It is a heterogeneous group of molecules because fatty acids with distinct lengths and degrees of saturation might form part of its structure; in this work, 1-oleyl-lysophosphatidic acid was the molecular species used. In addition to their metabolic roles, these lipid metabolites act as local intermediaries (diffusing through the intercellular media and serous fluids) and also as general messengers (circulating through the vascular system), and, for these reasons, they are considered “bioactive lipids”. Many of the most studied actions of LPA are mediated through a family of six G protein-coupled receptors (GPCRs) known as the LPA receptors (Choi et al., 2008). The subject of this work is the LPA receptor type 3 (LPA_3_ receptor), which triggers diverse signaling events such as phospholipase C activation, calcium mobilization, adenylyl cyclase inhibition, and stimulation of the mitogen-activated protein kinase pathway (Hama and Aoki, 2010; Solís et al., 2021). LPA_3_ receptors participate in many physiological events, such as embryo implantation, expression of antioxidant enzymes, decrease of apoptosis/survival promotion, and proliferation (Hama and Aoki, 2010; Solís et al., 2021). These receptors also play roles in pathological conditions such as neoplasias (Balijepalli et al., 2021), including ovarian cancer, a condition in which LPA_3_ expression is considered a poor prognosis marker and a therapeutic target (Zhao et al., 2022).

Recent work from our laboratory has shown that the ligands lysophosphatidic acid (LPA) and oleoyl-methoxy glycerophosphothionate (OMPT) exert similar actions but with marked potency and efficacy differences, which suggest that OMPT operates as a biased agonist on the LPA_3_ receptor (Solís et al., 2024c). Both agents increased equi-effectively LPA_3_ phosphorylation, but OMPT was more potent than LPA. OMPT’s efficacy was similar to that of LPA, but it was more potent to activate ERK 1/2. In contrast, OMPT was less effective but equipotent than LPA to increase intracellular calcium. Surprisingly, the EC_50_ values for LPA were 100-300 nM for all these distinct actions, whereas those for OMPT were ≈ 10 nM for some effects, but in rising intracellular calcium, it was similar to that of LPA (Solís et al., 2024c). These data suggest that the synthetic agonist might interact with multiple LPA_3_ receptor-binding sites. Agonist-induced receptor internalization also showed some differences; LPA- and OMPT-caused receptor internalization was fast, but the action of OMPT was much more marked. LPA-produced endocytosis was blocked by Pitstop 2 but not by Filipin, whereas that induced by OMPT was partially inhibited by both Pitstop 2 and Filipin and entirely by the combination of both (Solís et al., 2024c), again indicating differences in the actions of these agonists. In addition, when LPA-stimulated cells were rechallenged with 1 µM LPA, hardly any response was detected, i.e., a "refractory" state was induced (Solís et al., 2024c). However, a conspicuous reaction was observed when OMPT was the second stimulus. These data suggest that two binding sites for these agonists might exist in the LPA_3_ receptor, one showing a very high affinity for OMPT and another one, likely shared with LPA, with a lower affinity for both agonists (Solís et al., 2024c). Docking analysis evidenced that at least two agonist cavities exist in the LPA_3_ receptor: one located near the cytoplasmic face of this molecule (named Lower Cavity) and another one (Upper Cavity) situated near the receptor’s extracellular regions; sub-cavities within this large bindings pockets seem to exist (Solís et al., 2025).

Antidepressants are a large group of chemical agents used for the medical treatment of mood disorders, and many of these drugs are inhibitors of serotonin/ noradrenaline reuptake (O’Donnell and Shelton, 2011). It is disconcerting for both patients and psychiatrists that a very long treatment is required to evidence any clinical improvement (4 to 8 weeks on average). Therefore, the possibility of other mechanisms of action (for example, modulation of genetic expression) and the role of targets distinct from the serotonin/ catecholamine sodium-dependent transporters has been considered (O’Donnell and Shelton, 2011). Interestingly, the pioneering work of Olianas et al. (Olianas et al., 2015) showed that different classes of antidepressants could induce insulin-like growth factor-I receptor transactivation, stimulation of ERK1/2 signaling, and cell proliferation in CHO-K1 fibroblasts and that these effects were mediated through LPA_1_ receptor stimulation. This antidepressant effect has now been observed in various experimental models (Banks et al., 2018; Kajitani et al., 2016; Kajitani et al., 2024; Olianas et al., 2015, 2017, 2019, 2020, 2023). Recent evidence indicates that the ability of these agents to activate LPA receptors is not restricted to the LPA_1_ subtype but common to other members of this family, including the LPA_3_ subtype (Kajitani et al., 2024; Olianas et al., 2023).

Consequently, we explored whether some of these drugs (imipramine, amitriptyline, and paroxetine; Supplementary Fig. S1) exert their effects by interacting with the LPA_3_ binding cavities, both experimentally, in cultured cells, and *in silico* through docking studies. Two of these antidepressants, imipramine and amitriptyline, have a very similar chemical structure belonging to the classic “tricyclic antidepressants” (O’Donnell and Shelton, 2011). In contrast, paroxetine is a member of the “selective serotonin uptake inhibitors” group (O’Donnell and Shelton, 2011), with a different chemical configuration, i.e., it is a piperidine containing benzodioxole (Supplementary Fig. S1). Our data clearly show that antidepressant activation of LPA_3_ receptors takes place through interaction with the Upper Cavity. Despite this similarity, marked differences were observed in the effects of these agents and in the structural elements that participate in their chemical interrelation with the receptor’s Upper Cavity.

## 2. Material and Methods

### Reagents

1-Oleyl lysophosphatidic acid (LPA) and 2S-(1-oleoyl-2-O-methyl-glycerophosphothionate) (OMPT) were from Cayman Chemical Co. (Ann Arbor, MI, USA). Imipramine hydrochloride, amitriptyline hydrochloride, and paroxetine were generous gifts from Psicopharma S. A. de C.V. (Mexico) (http://www.psicofarma.com.mx/). Dulbecco’s modified Eagle’s medium, antibiotics, and Fura-2 AM were purchased from Invitrogen-Life Technologies (Carlsbad, CA, USA). The LPA_3_ receptor sequence was fused at the carboxyl terminus with the green fluorescent protein and cloned into the pCDNA5/FRT/TO plasmid (Bioinnovatise, Inc., Rockville, MD, USA) to employ the inducible HEK 293 Flp-In TREx expression system (Ward et al., 2011). The source and catalog number of cells and antibodies and data for other materials are indicated in our previous publications (Solís et al., 2024a, b; Solís et al., 2024c).

### Intracellular calcium concentration

Determinations were performed as previously described (Solís et al., 2024a). In brief, the cells were serum-starved and treated with doxycycline hyclate to induce LPA_3_ expression, loaded with 2.5 µM Fura-2 AM for 1 h. and later carefully detached from the Petri dishes, then washed to eliminate unincorporated dye, and maintained in suspension (García-Sáinz et al., 2010). In the present work, very low (titrated to avoid loss of responsiveness) concentrations of trypsin were used to detach the cells, which resulted in better cell yields and a more extensive response to OMPT. Two excitation wavelengths (340 and 380 nm) and the emission wavelength of 510 nm were employed. Intracellular calcium levels were calculated as described by Grynkiewicz et al. (Grynkiewicz et al., 1985). In a series of experiments, cells were subjected to two sequential stimuli to determine if they could respond again; the second stimulus was added without removing the first one, as described (Solís et al., 2024c).

### ERK 1/2 phosphorylation

Cells were serum-starved for 4 h and then activated for the indicated time with the agents to be tested; after this incubation, cells were washed twice and lysed (Solís et al., 2024a); the lysates were centrifuged, and proteins contained in supernatants were denatured with Laemmli sample buffer (Laemmli, 1970) and separated by SDS-polyacrylamide gel electrophoresis. Proteins were electrotransferred onto membranes, and immunoblotting was performed. Total- and phospho-ERK 1/2 levels were determined in the same membranes for each experiment; the baseline value was considered 100% for normalization. In all experiments, cell incubations with LPA were performed in parallel; the cell extracts were also run together to properly compare the effects of the distinct agents with those of LPA. Also, in a series of experiments, cells were subjected to two sequential stimuli to determine if they could respond again; the second stimulus was added without removing the first one, as performed in the intracellular calcium experiments.

### Receptor internalization and membrane fluorescence

Cells were seeded on glass-bottomed Petri dishes for 12 h, and LPA_3_ receptor expression was induced with doxycycline (Solís et al., 2024a). Before the experiment, the cells were serum-fasted for 1 h and then stimulated for the times and with the agents indicated. Cells were washed and fixed as described (Solís et al., 2024a). The green fluorescent protein was excited at 488 nm and emitted fluorescence registered at 515-540 nm. The plasma membrane was delineated using the differential interference contrast images to determine receptor internalization. Each cell’s intracellular fluorescence (i.e., excluding the plasma membrane) was quantified as “integrated density”, employing the ImageJ software (Rasband, 1997-2004), as described (Solís et al., 2024a). The baseline intracellular/total fluorescent ratio was considered 100% (the S.E.M. of the different samples in each experiment indicated the internal variation). Plasma membrane fluorescence [1-(intracellular/total fluorescent ratio)] was determined; the baseline value was considered 100 %. Usually, 10-14 images were taken from 3 or 4 cultures obtained on different days for each condition.

### Video experiments

The video experiments employed a confocal Zeiss LSM800 microscope with a temperature- and atmosphere (CO_2_ and humidity)-controlled chamber as described (Solís et al., 2024a). Excitation, emission, and recording details were as previously described (Solís et al., 2024a, b). It should be considered that cells and organelles move (i.e., migrate and change their form) during the experiments, entering and leaving the observation plane and that LPA_3_ stimulation induces cell contraction and migration (Solís et al., 2024a, b).

### Docking studies

The LPA_3_ receptor has not been crystallized yet. Based on the amino acid sequence obtained from the Uniprot database using the Q9UBY5 entry (https://www.uniprot.org/uniprotkb/Q9UBY5/entry), the most likely 3D structure was predicted employing the SWISS-MODEL server (https://swissmodel.expasy.org/) (Solís et al., 2025). A protein 3D structure refinement to reach a structurally relaxed conformation was executed to perform accurate docking simulations using ReFOLD (https://www.reading.ac.uk/bioinf/ReFOLD/ReFOLD3_form.html). The refined protein with the best score was used for the docking study to explore receptor cavities capable of agonist recognition (Solís et al., 2025). Possible ligand identification pockets were explored using the BetaCavityWeb (http://voronoi.hanyang.ac.kr/betacavityweb/) and the Fpocketweb (http://durrantlab.com/fpocketweb) servers. The 3D configurations of the agents studied were obtained from PubChem (https://pubchem.ncbi.nlm.nih.gov/) (ID 5497152 for LPA and ID 16078994 for OMPT (as reported in (Solís et al., 2025)) and molecular structure minimization was carried out using the Avogadro program (https://avogadro.cc/docs/tools/auto-optimize-tool/); the lengths and angles of the links and junctions of atomic interactions as well as their geometry were appropriated with the optimized energy under the mentioned structure minimization The antidepressant information as 3D was obtained from the Protein Data Bank’s RCSB archive (https://www.rcsb.org/) (ID 2q72 for imipramine, ID 3apv for amitriptyline, and ID 4mm4 for paroxetine). All simulations were performed using the LINUX operating system (Centus) and the Lamarckian genetic algorithm for scoring sampling using Autodock, as described (Solís et al., 2025). The docking simulations had a maximum number of 1 x 10^7^ energy evaluations and a population of 100 randomized individuals (represented by the 100 best evaluations) (Solís et al., 2025). The 2D maps showing the receptor’s residues interacting with the ligands were obtained using the Discovery Studio Visualizer (https://discover.3ds.com/discovery-studio-visualizer-download). The thermodynamic estimations of the ΔG and Kd values were obtained directly from the “*.dlg files” after docking simulations using the program Autodock tools 1.5.6 (ADT) (https://autodocksuite.scripps.edu/adt/).

### Statistical analyses

The data are presented as the means ± standard errors of the means with the number of samples indicated. Statistical comparison between two groups was performed using the Student’s t-test, and when more groups were compared, the ordinary one-way ANOVA with the Bonferroni post-test was employed. These analyses were performed using the software included in the GraphPad Prism program (version 10.2.2).

## 3. Results

### Intracellular calcium

As shown in Fig. 1, the natural agonist, LPA, was the most efficacious, markedly increasing intracellular calcium in a concentration-dependent fashion (EC_50_ ≈ 60 nM). As anticipated, the OMPT effect was smaller but showed a similar apparent potency (EC_50_ ≈ 60 nM). Paroxetine had an efficacy similar to OMPT, but the EC_50_ value was considerably higher (i.e., ≈ 30 µM). Amitriptyline and imipramine increased intracellular calcium to a minimal extent, and no saturation was apparent within the range of concentrations tested. The findings with the tricyclic antidepressants show similarity with what has been observed in other cellular models (Kajitani et al., 2024; Olianas et al., 2023) and indicate that the three distinct drugs tested were low-affinity LPA_3_ receptor agonists with lesser efficacy than LPA). Based on these findings, in all the following experiments, the concentration of the ligands was 1 µM for LPA and OMPT and 30 µM for the antidepressants.

**Fig. 1.**
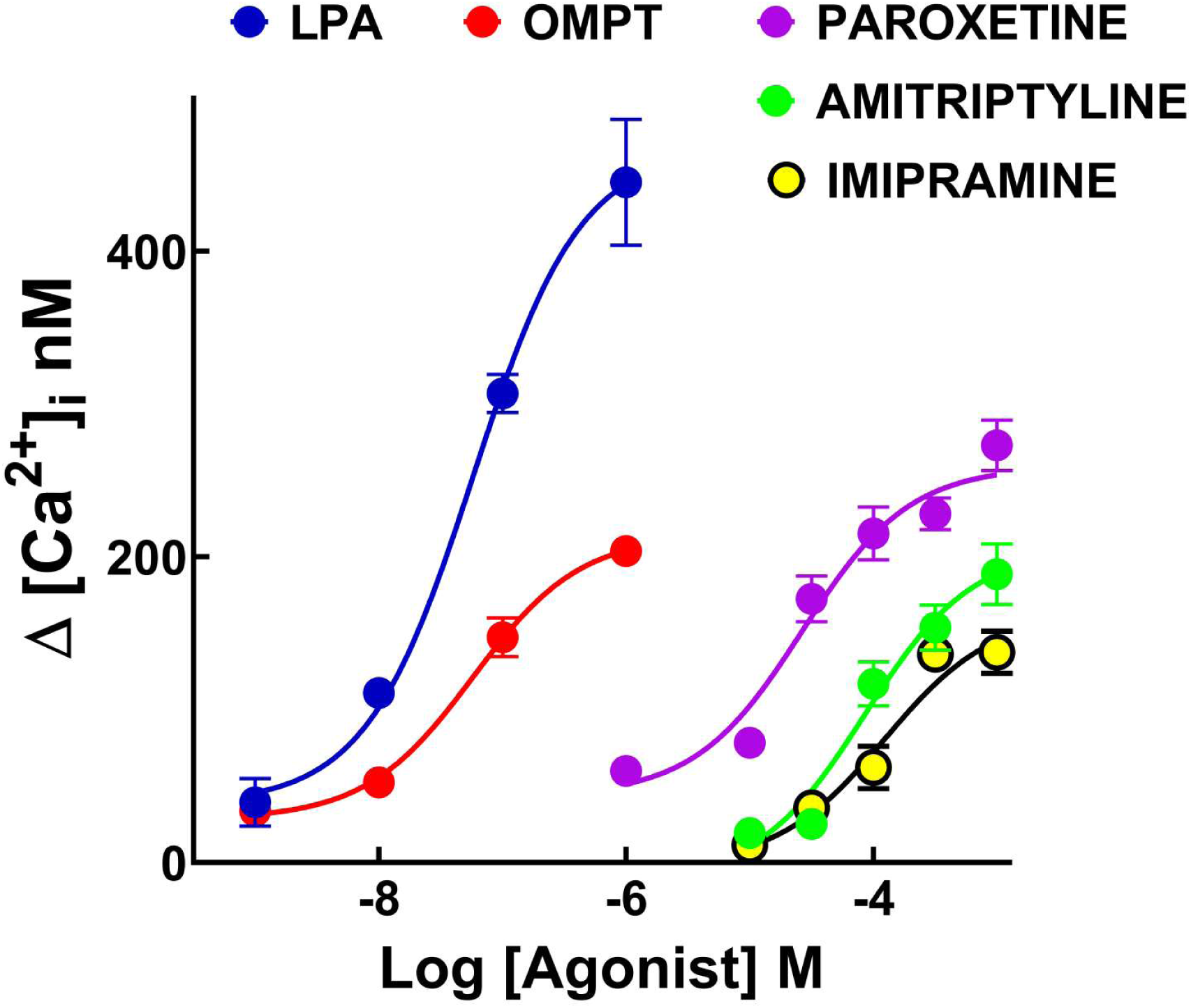
Increases in intracellular calcium in response to different concentrations of LPA (blue symbols and line), OMPT (red symbols and line), paroxetine (purple symbols and line), amitriptyline (green symbols and line), and imipramine (yellow symbols with black borders and black line). The means are plotted, and vertical lines indicate the SEM of 6-8 experiments performed with distinct cell cultures.

In the first group of columns of Fig. 2, the effects of LPA, OMPT, paroxetine, imipramine, and amitriptyline (30 µM) are depicted, and the response magnitude ratios (agent/LPA) are indicated above each bar. OMPT and paroxetine induced, at the concentrations tested, an action corresponding to ≈ 60% of that of LPA, whereas imipramine and amitriptyline were much less effective (≈ 30% of LPA action) (Supplementary Fig. S2 presents representative calcium tracings). The second group of columns shows what happens when cells are stimulated with LPA and rechallenged with the distinct agents without removing the added LPA. Cells redefied with LPA exhibited a minimal (marginal) response (≈ 10% of the initial LPA effect), confirming that the LPA_3_ receptor becomes “refractory” (or selectively desensitized) (Solís et al., 2024c) to a new LPA stimulation (Fig. 2, Supplementary Fig. S3, panel A). Under the same conditions, OMPT and paroxetine induced relatively small but conspicuous responses (Fig. 2, Supplementary Fig. S3 panels B and C). Interestingly, rechallenging cells with imipramine or amitriptyline decreased intracellular calcium even below the initial (time 0’) baseline values (Fig. 2, Supplementary Fig. S3 panels D and E).

**Fig. 2.**
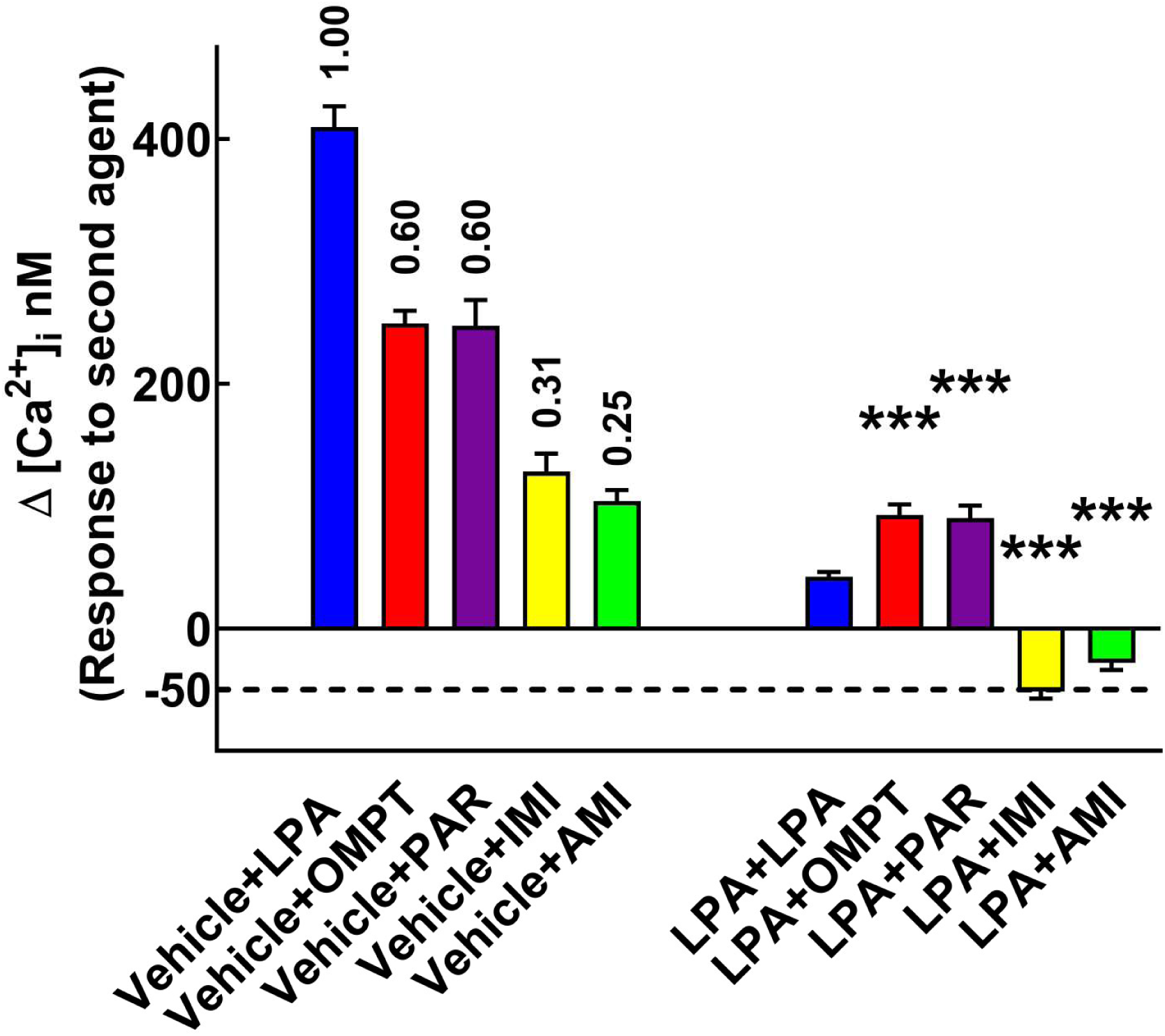
Cells were preincubated with vehicle (left group of columns) or 1 µM LPA (right group of columns) for 100 seconds. After this preincubation, cells were rechallenged with 1 µM LPA (blue bars), 1 µM OMPT (red bars), 30 µM paroxetine (purple bars), 30 µM amitriptyline (green bars), or 30 µM imipramine (yellow bars with black borders). The increases in intracellular calcium concentration in response to the second stimulus were determined; means are plotted, and vertical lines indicate the SEM of 5-8 determinations using distinct cell cultures. *** p < 0.001 vs. the LPA+LPA group. The numbers above the left group of bars are the activities of the agents relative to LPA.

### ERK 1/2 phosphorylation

We next examined this longer-term reaction to these agents. Fig. 3 shows the concentration-response curves for the different drugs studied (panel A) and the time courses of ERK 1/2 phosphorylation (panel B). As anticipated, LPA and OMPT induced rapid and robust effects on this parameter (Solís et al., 2024c). Imipramine produced a swift and vigorous reaction but only at the highest concentrations tested (30 and 100 µM (Fig. 3). Surprisingly, amitriptyline and paroxetine were unable to generate any fast and consistent increase in ERK 1/2 phosphorylation and at more extended times of incubation these agents even decreased this parameter below the baseline (Fig. 3B). Representative Western blots are presented in Supplementary Fig. S4.

**Fig. 3.**
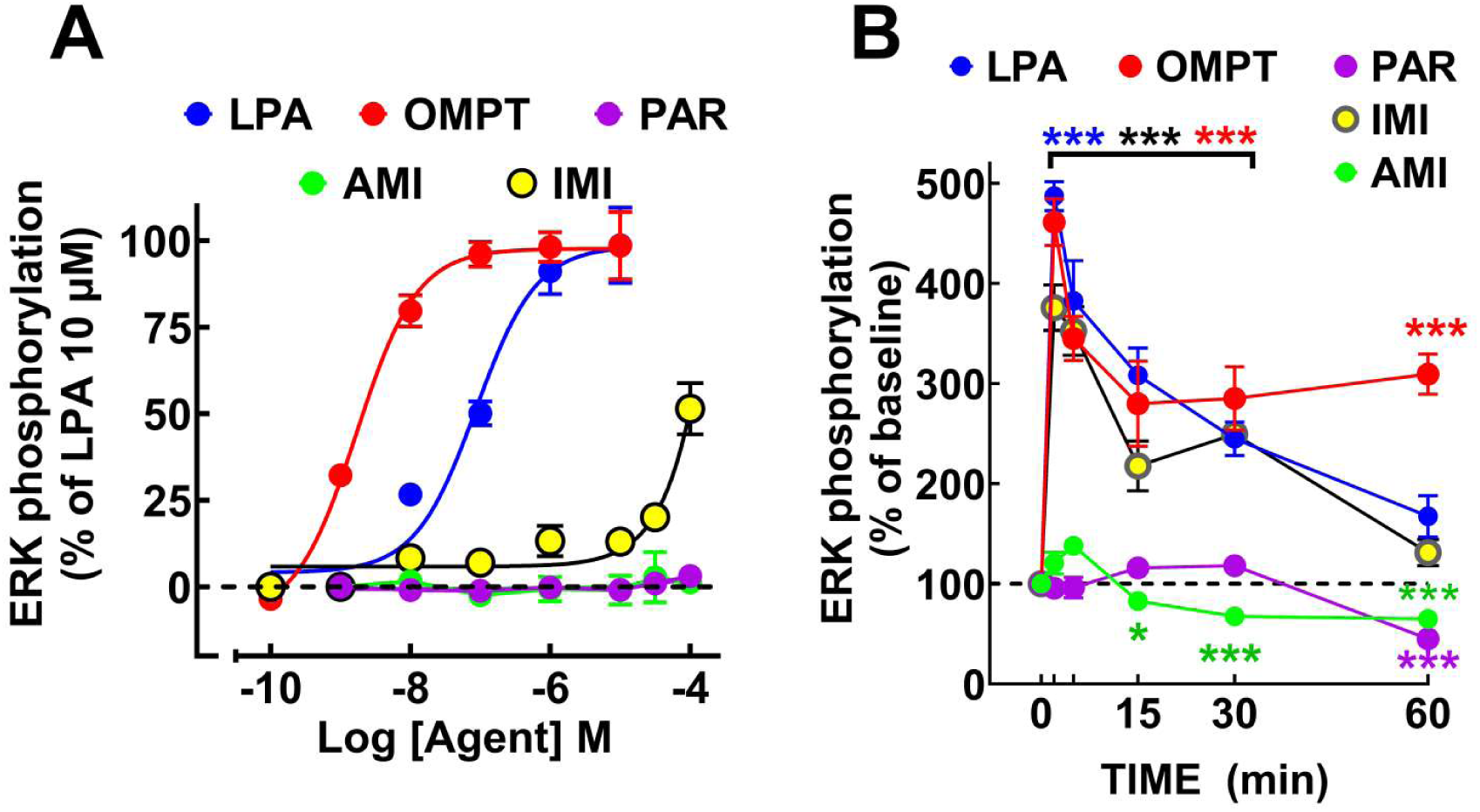
Effect of LPA, OMPT, and antidepressants on ERK 1/2 phosphorylation. In panel A, cells were incubated for 2 min with the indicated concentrations of LPA (blue symbols and line), OMPT (red symbols and line), paroxetine (purple symbols and line), amitriptyline (green symbol and line), or (yellow symbol with black borders and line). In panel B, cells were incubated for the times indicated with 1 µM LPA (blue symbols and line), 1 µM OMPT (red symbols and line), 30 µM paroxetine (purple symbols and line), 30 µM amitriptyline (green symbol and line), or 30 µM imipramine (yellow symbol with black borders and line). The means are plotted, and vertical lines indicate the SEM of 4-6 determinations using distinct cell cultures. *** p < 0.001 and * p < 0.05 vs. the respective baseline (time 0’) value, color-coded.

Next, we examined if cells incubated with LPA for 60 min could respond to the addition of the distinct drugs, as performed for the intracellular calcium determinations described above. Contrary to what we observed in the calcium experiments, LPA-treated cells rechallenged with LPA responded rapidly, although the effect was reduced to approximately 50% of the initial one (Fig 4). The data indicate that this action does not become "refractory" to a second stimulus by LPA. The experimental conditions of these two types of experiments are very different (the course of the response or the incubation times, among others). When LPA-activated cells were challenged with OMPT or imipramine, rapid and robust reactions similar to that of LPA were detected (Fig. 4). Interestingly, amitriptyline and paroxetine had an opposite effect, i.e., these antidepressants decreased ERK 1/2 phosphorylation even below the value recorded before the initial LPA stimulation; such reductions were statistically significant and the action of paroxetine was evident as soon as 5 min after its addition to the cells (Fig. 4). Representative Western blots are presented in Supplementary Fig. S5). It is worth mentioning that all of these agents induce marked changes in cell shape (contraction), and in the case of the cells rechallenged with the tricyclic antidepressants, amitriptyline, and imipramine, we detected that cell detachment took place, particularly at 30 and 60 min of the second stimulation. Consequently, when these antidepressants were used as a second stimulus, the medium was recovered (indicated as AMI R (for Recovered) and IMI R in Fig. 5), centrifuged, and the pellet (detached cells) was added to the cell extract to appropriately compare the results (i.e., obtaining similar total ERK detection; comparative (Recovered vs. not recovered) pERK and ERK Western blots are presented in Supplementary Fig. S6).

**Fig. 4.**
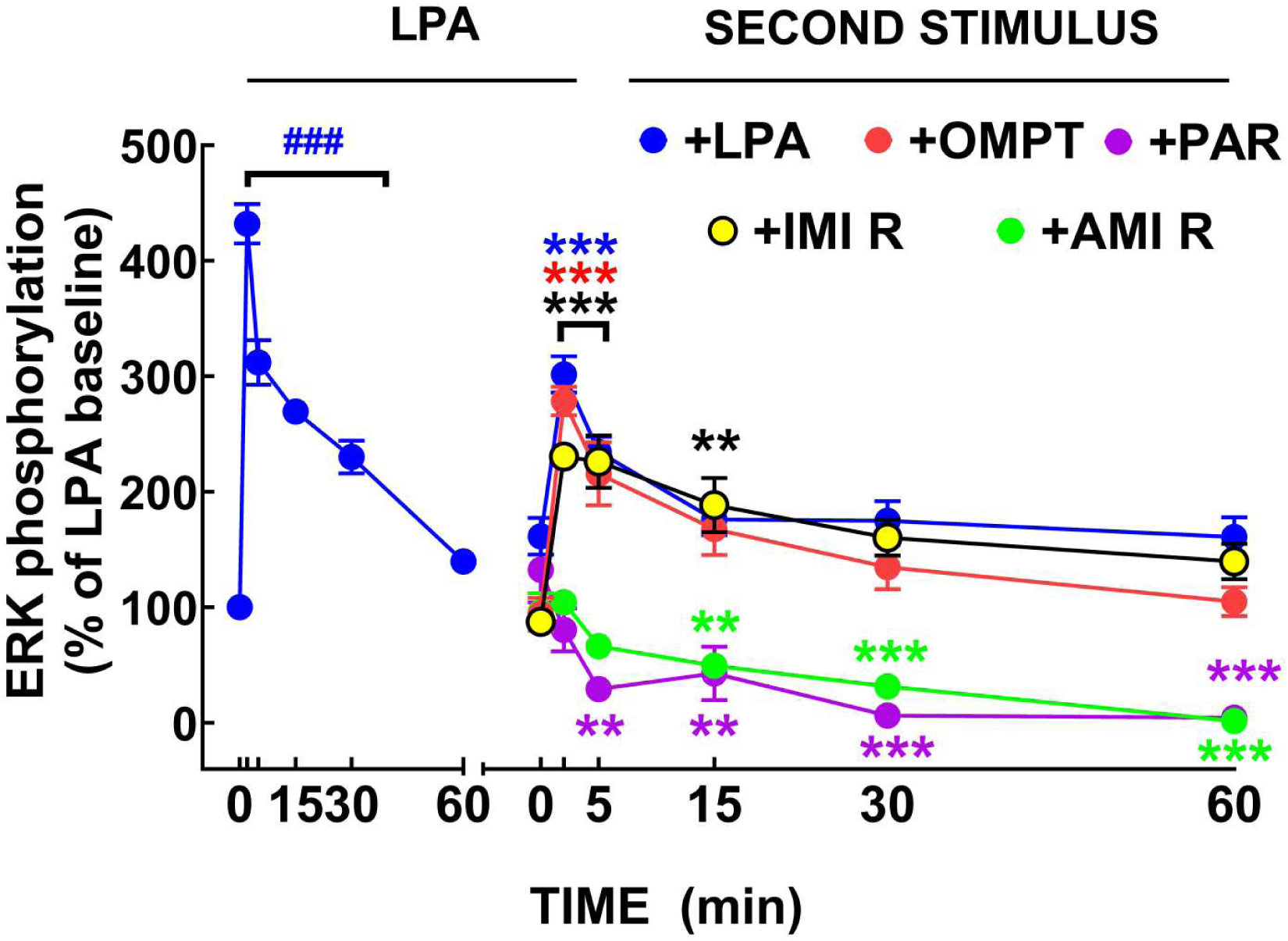
Changes in ERK 1/2 phosphorylation in cells incubated with 1 µM LPA for 60 min. Following this incubation, cells were rechallenged with 1 µM LPA (blue symbols and line), 1 µM OMPT (red symbols and line), 30 µM paroxetine (purple symbols and line), 30 µM amitriptyline (green symbols and line), or 30 µM imipramine (yellow symbols with black borders and black line). The means are plotted, and vertical lines indicate the SEM of 5-8 determinations using distinct cell cultures. ### p < 0.001 vs. initial baseline (time 0’) value.*** p < 0.001 and ** p < 0.01 vs. respective second stimulus time 0’ values; color coded.

**Fig. 5.**
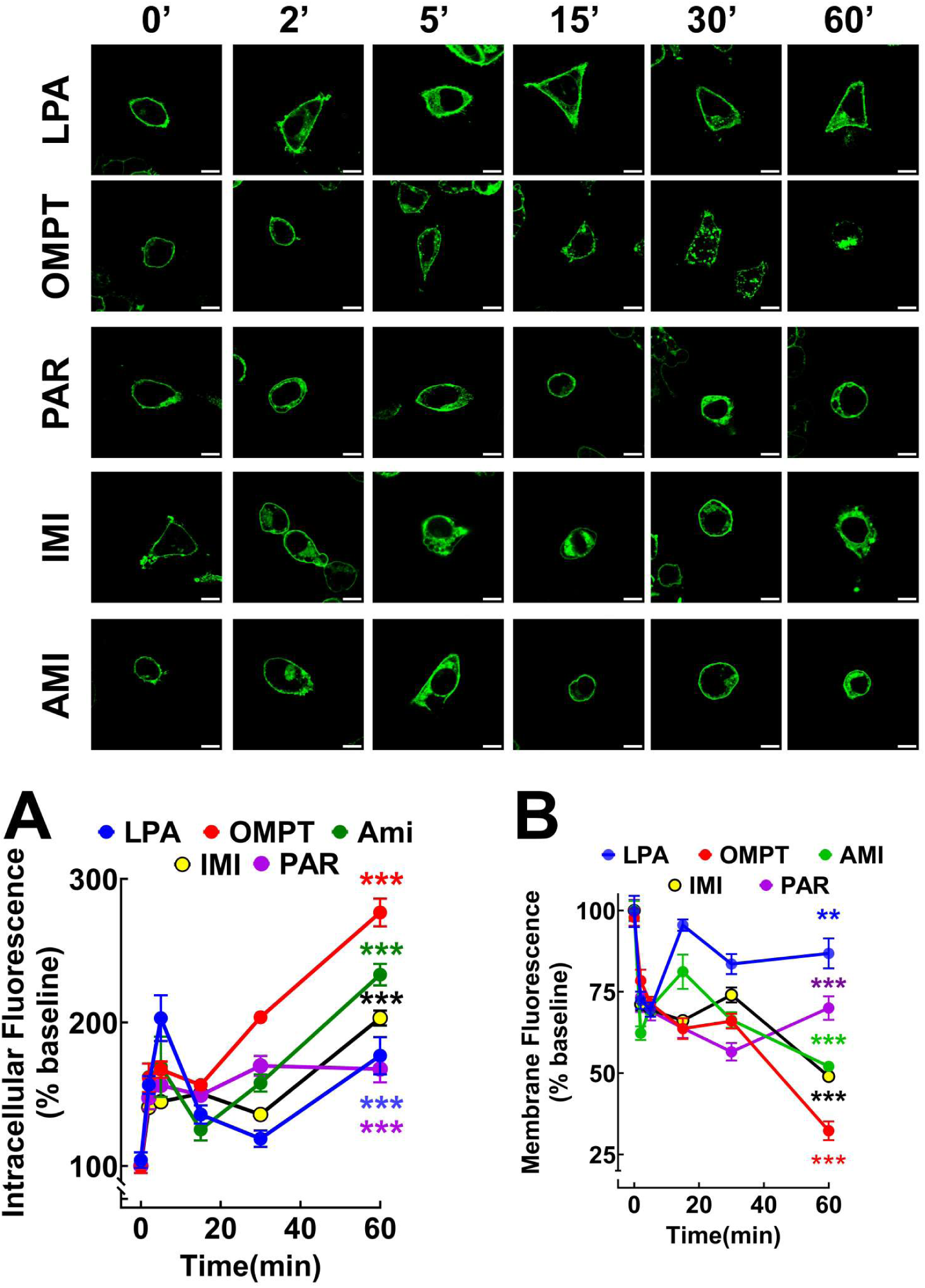
Intracellular (panel A) and membrane (panel B) fluorescence of cells incubated with 1 µM LPA (blue symbols and line), 1 µM OMPT (red symbols and line), 30 µM paroxetine (purple symbols and line), 30 µM amitriptyline (green symbols and line), or 30 µM imipramine (yellow symbols with black borders and black line). The means are plotted, and vertical lines indicate the SEM of 14 determination of each of the 4 experiments performed using distinct cell cultures. *** p < 0.001 vs. respective time 0’ values; color coded.

### Receptor internalization and plasma membrane fluorescence

As reported previously, we used the accumulation of intracellular fluorescence as an index of LPA_3_ internalization (Solís et al., 2024a, b; Solís et al., 2024c). As expected, LPA triggered a rapid LPA_3_ receptor internalization, reaching its maximum at 5 min and decreasing later. In contrast, OMPT induced a similar initial swift receptor endocytosis followed by a second slower increase of intracellular accumulation of fluorescence maintained until the end of the incubation (Fig. 5A) (Solís et al., 2024c). Imipramine, amitriptyline, and paroxetine triggered a similar pattern of internalization as OMPT, although of lesser magnitude (Fig. 6). We also determined the change in plasma membrane fluorescence. We previously showed that LPA only marginally decreases plasma membrane LPA_3_-green fluorescent protein signal in this cellular model, but, in contrast, OMPT induces a marked reduction in fluorescence (Solís et al., 2024c). This finding was confirmed here, and it was observed that the antidepressants also decrease plasma membrane fluorescence. However, it was less pronounced than that triggered by OMPT (Fig. 5B). The effects of these drugs evidenced marked differences as shown in the videos (in all videos, format A exhibits the fluorescence images and format B, the fluorescence with the differential interference contrast overlapped images). In Videos 1A and 1B, the action of paroxetine is depicted; it can be observed that after the addition of this agent, the fluorescence that finely defines the plasma membrane rapidly aggregates, forming patches in such a way that rather than being delimited by a continuous fluorescent line, a “dotted contour” pattern marked it. Furthermore, we observed blebbing and the appearance of a large amount of “optically dense vesicles” or granules (some of them fluorescent, i.e., likely containing receptors) located abundantly around the cells (we wonder if some attachment to the cells remained). An abrupt interruption of the streaming of cytoplasm within the cells (cyclosis) was also evidenced.

**Fig. 6.**
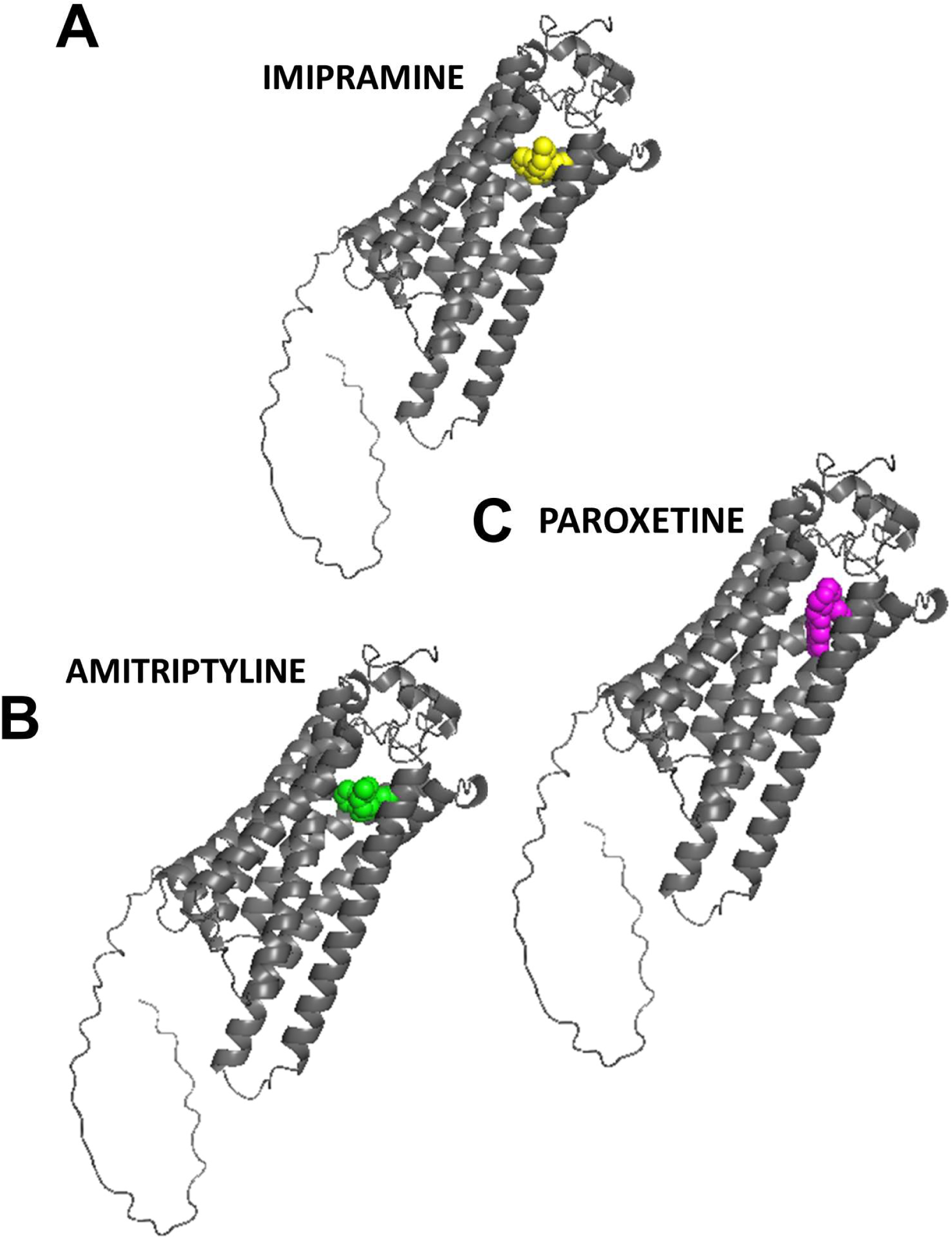
3D Graphical representation of the docking of the refined LPA_3_ receptor to the ligands: imipramine (yellow, panel A), amitriptyline (green, panel B), and paroxetine (purple, panel C).

The morphological actions of amitriptyline (Videos 2A and 2B) and imipramine (Videos 3A and 3B) were also impressive; i.e., after the addition of these tricyclic antidepressants, the cells contracted, and a very active blebbing took place, forming enormous vesicles protruding from the plasma membrane that, in some cases, appear to dissociate from the cells and, in others, seem to join them. Amitriptyline treatment also induced some accumulation of dense granules around the cells; the effect was much less intense and slower than that of paroxetine. We were puzzled by these findings and tested the reaction to these antidepressants in uninduced cells (i.e., not expressing the LPA_3_ receptors), which responded to paroxetine addition by blebbing and an abrupt blockade of cyclosis; however, no presence of dense granules was observed (Video 4). Similarly, in uninduced cells, imipramine and amitriptyline triggered contraction and blistering with the formation of enormous vesicles (Videos 5 and 6), although it appeared to be less intense than in LPA_3_-expressing cells (attempts to quantify these were unsuccessful).

### Docking simulations

We recently described an LPA_3_ receptor 3D structure based on the amino acid sequence and how it was refined and validated (Solís et al., 2025). The molecular docking simulation showed that the LPA_3_ receptor interacting sites for LPA and OMPT differed and varied with the docking conditions employed (Solís et al., 2025). LPA can associate with the receptor via the Intracellular Loops 2 and 3 with what we denominated Lower Cavity, very close to the cytoplasm, but also with an Upper Cavity with the receptor’s amino acids in transmembrane regions 5, 6, and 7, and the extracellular loop 3 (Supplementary Fig. S7). In contrast, OMPT interacted mainly with the Upper Cavity (Solís et al., 2025) (Supplementary Fig. S7). When the docking simulations were focused on Trp102, both LPA and OMPT were found in the Upper Cavity (Solís et al., 2025) (Supplementary Fig. S7).

The three antidepressants studied occupied domains within the Upper Cavity (Fig. 6, panels A-C). The use of the same cavity is further illustrated in Supplementary Fig. S7; it can be observed that these drugs bind to sites within this recognition pocket. However, the ligand-receptor interactions are not structurally identical. Imipramine and amitriptyline are associated with the transmembrane helix 3, 5, 6, and 7 but in a different structural orientation. The configuration of these two tricyclic antidepressants is very similar, and both interrelate analogously with the Upper Cavity (i.e., like “hanging from vines”). Paroxetine seems to be associated mainly with transmembrane helixes 3 and 6 (i.e., in a “climbing tree” way) (Fig. 6, panel C). A 3D image (PyMOL) of the three antidepressants simultaneously interacting with the LPA_3_ receptor is presented in Fig. 7. The amino acids that interrelate with each of the drugs and the thermodynamic and affinity parameters (ΔG and Kd), obtained with the "*.dlg files" using Autodock tools 1.5.6, are presented in Table 1. Supplementary Fig. S8 shows cartoons of the LPA_3_ receptor indicating in color the interacting amino acids with each antidepressant. Similarly, the 2D maps of the ligand-receptor interrelations obtained from the Discovery Studio program are depicted in Supplementary Fig. S9. Many of the LPA_3_ residues these drugs associate with are shared (Table 1). Of the 12 amino acids that are sites of contact with imipramine, 9 of them also interact with amitriptyline and 7 with paroxetine; Tyr183 is a common point of connection for amitriptyline and paroxetine but not for imipramine (Table 1). A total of 6 residues, i.e., Trp152, Asp110, Gln106, Leu259, Phe278, and Leu279, are common interaction sites for the three antidepressants studied. The possibility that some of these amino acids could also be shared with the agonist, LPA, and OMPT was analyzed by the docking studies focused on Trp102 with distinct ligand charges. Interestingly, when the ligand charge was -2 or 0, only Leu279 was a common point of interaction for all the antidepressants, and when the agonist charge was -1, the degree of agent-receptor association was much more important. LPA, imipramine, amitriptyline, and paroxetine shared interaction with Gln106 and Leu279; LPA, imipramine, and amitriptyline contacted Leu109; and Lys275 binds to LPA and paroxetine.

**Fig. 7.**
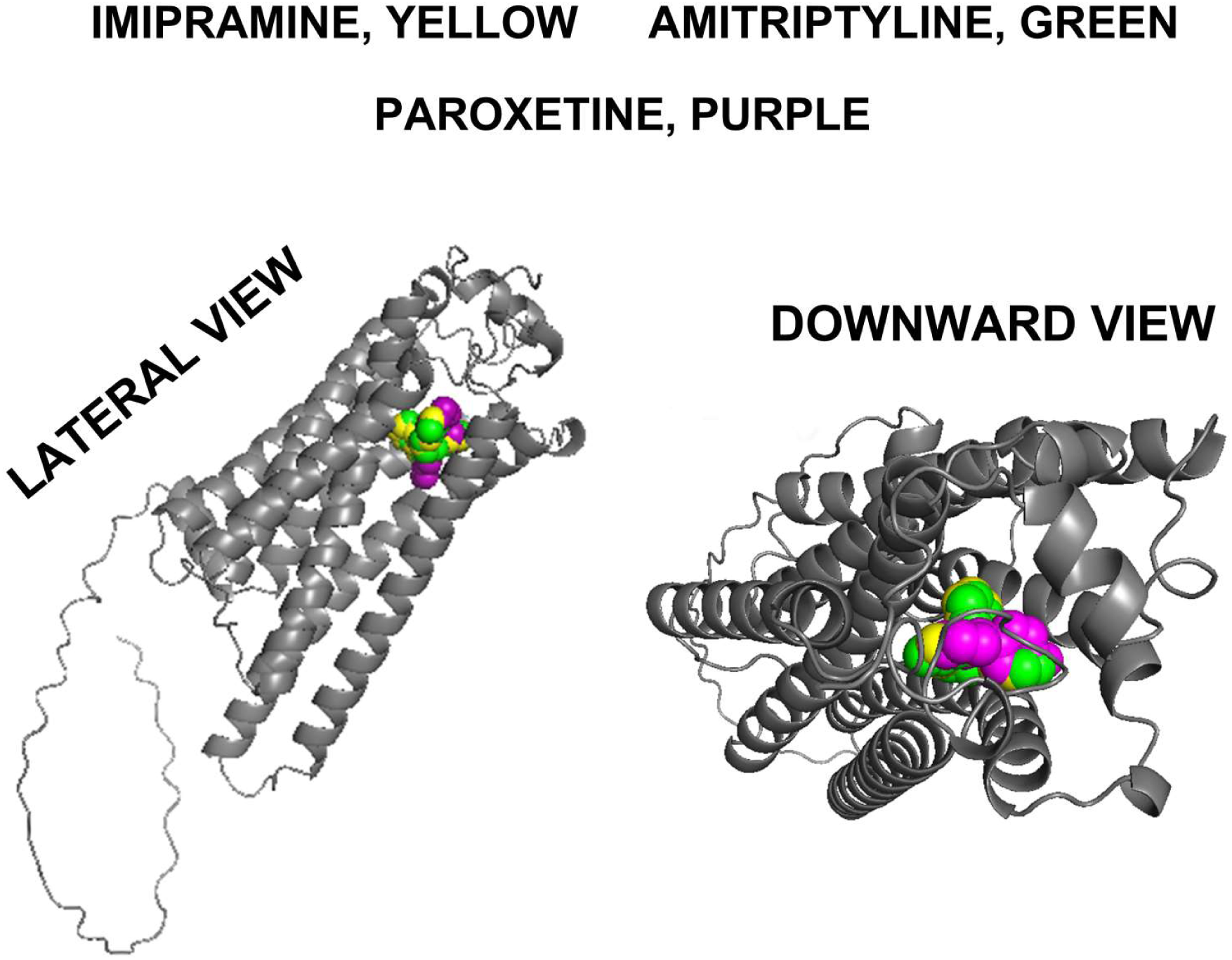
3D Images of the LPA_3_ receptor refined model docking with imipramine (yellow), amitriptyline (green), and paroxetine (purple), obtained using PyMOL. Lateral and downward views are presented.

**TABLE 1.** Interaction of distinct ligands with the LPA_3_ receptor. Thermodynamic parameters and amino acids involved.

| **Ligand** | **$\Delta G$**<br>**(kcal/mol)** | **K<sub>d</sub>**<br>**(nM)** | **Interactions** |
| --- | --- | --- | --- |
| **Imipramine** | -8.76 | 380 | Trp252, Ala282, Asp110, Trp191, Leu109, Leu256, Gly255, Gln106, Leu259, Val258, Phe278, Leu279 |
| **Amitriptyline** | -8.01 | 1340 | Trp252, Asp110, Leu109, Ala282, Trp191, Tyr183, Gln106, Leu259, Phe278, Leu279 |
| **Paroxetine** | -8.49 | 595 | Trp252, Cys251, Asp110, Gly255, Phe278, Leu259, Tyr183, Gln106, Ala180, Lys275, Leu279 |

OMP had in common with antidepressants more contact sites in the LPA_3_ receptor. i.e., Trp252, Ala282, Asp110, Trp191, Leu256, Gln106, Phe278, Leu279, Tyr183, and Lys275. A 3D image (Pymol) of the three antidepressants, LPA and OMPT, interacting with the LPA_3_ receptor is presented in Supplementary Fig. S10 employing the Trp102-focused data with the distinct agonist charges. The images show that under the different charge conditions, the position of LPA does not overlap with that of the antidepressants; with charges 0 and -2, there seems to exist proximity between LPA and OMPT but not with the drugs. In the image obtained using LPA and OMPT charge -1, OMPT appears to interact with the receptor in a position closer to that of the antidepressants. These data suggest that the ligands might interplay with different sub-cavities within the Upper Cavity and that such distinct associations could be related to the diverse functional effects observed (Supplementary Fig. S10).

## 4. Discussion

It has been noticed that antidepressants exhibit activity on LPA receptors, particularly LPA_1_ (Kajitani et al., 2016; Olianas et al., 2015, 2017, 2019, 2020), but it was recently reported that they also stimulate the LPA_3_ subtype (Kajitani et al., 2024; Olianas et al., 2023). The findings that the ligands LPA and OMPT showed differences in their cellular actions and bound to distinct sites in the LPA_3_ receptor (Solís et al., 2025; Solís et al., 2024a, b; Solís et al., 2024c) prompted us to examine other reported agonists. Among these agents are the antidepressants imipramine, amitriptyline, and paroxetine, studied in this work. A significant contribution of this research is to provide evidence that the effects of these drugs differ from those of LPA and resemble, to some extent, the repercussion of OMPT on calcium signaling *in cellulo* and that they interact with the LPA_3_ receptor Upper Cavity. It is known that docking simulations generally match the experimental evidence, as has been observed with different targets (Shen et al., 2023; Zhao et al., 2024). Our studies evidenced that the ligands recognize the LPA_3_ receptor with favorable free energy values; however, despite the reliability of the binding affinities predicted by docking simulations, experimental validation is recommended, as was achieved in this work.

Another important aspect of this research is that, although the antidepressants studied interact with the LPA_3_ Upper Cavity, they did not associate with identical residues and were found in distinct angular positions in relationship with the receptor vertical axis. It is worth noticing that the antidepressants do not trigger the same cellular effects, which allows us to suggest that the receptor’s structural movements induced by the receptor-ligand association could differ in each case, resulting in distinct configurations that might affect its interaction with the elements involved in signaling, like G proteins, β-arrestins, and others. Such putative conformational changes possibly define the varied antidepressant actions observed when compared to LPA and also among them. This interpretation requires confirmation by molecular dynamics and validation through structural studies (X-ray crystallography or cryo-electron microscopy), presently unreachable goals for us.

Several parameters examined in our study have not been reported before, to the extent of our knowledge; one of them was the ability of these agents to increase intracellular calcium. Considering that LPA is a full agonist, OMPT and paroxetine are partial agonists and imipramine and amitriptyline exhibited relatively low efficacy, it has been observed that paroxetine is hardly capable of stimulating LPA_1_ receptors (Kajitani et al., 2016); however, in contrast, it shows a general G protein activating effect (evidenced by secondary TGFα shedding) in cells expressing LPA_3_ receptors (Kajitani et al., 2024), which is consistent with our findings. In the case of paroxetine, the noticeable differences in efficacy increasing intracellular calcium and ERK phosphorylation indicated a marked bias toward one action over another (biased agonism).

As anticipated, in the rechallenge experiments using LPA-stimulated cells, there was hardly any LPA-induced increase in intracellular calcium (Solís et al., 2024c). OMPT and paroxetine triggered reduced but distinguishable effects, and the tricyclic antidepressants decreased intracelllular calcium concentration even below initial baseline levels (i.e., under these conditions, imipramine and amitriptyline functioned as antagonists or inverse agonists). This marked reduction in intracellular calcium was not too surprising since it has been reported that the LPA_3_ receptor exhibits constitutive activity (blocked by Ki1645) (Solís et al., 2024c); however, it was puzzling that imipramine and amitriptyline showed distinct profiles alone (weak agonists) compared to when in the presence of LPA (inverse agonists).

When ERK 1/2 phosphorylation was studied, paroxetine induced no stimulation, indicating that this antidepressant is noticeably biased toward calcium signaling; it also markedly decreases ERK 1/2 phosphorylation in LPA-treated cells. Imipramine showed an opposite profile; it effectively activates ERK 1/2 phosphorylation, while its action on the intracellular calcium concentration was small. The tricyclic agents, amitriptyline and imipramine, exhibited similar effects on calcium signaling, i.e., these drugs were partial agonists with minimal intrinsic activity by themselves and behaved as inverse agonists in LPA-stimulated cells. Surprisingly, when ERK 1/2 phosphorylation was studied, these structurally related agents elicited distinct responses, i.e., imipramine was an efficacious partial agonist, while amitriptyline did not show any significant effect. In LPA-stimulated cells, imipramine, as a second stimulus, increased ERK 1/2 phosphorylation, whereas amitriptyline was an inverse agonist/ antagonist. These data indicate that, although these structurally similar antidepressants interact with the same LPA_3_ Cavity, they induce distinct functional responses. Caution should be exercised to avoid generalizations.

The receptor internalization studies showed that all the agents examined could induce this process, although with different kinetics and magnitudes. Similarly, fluorescence analysis of the plasma membrane indicated that OMPT was much more effective than LPA or the three distinct antidepressants examined in decreasing the density of the receptor. However, the confocal video studies revealed that these drugs triggered very complex processes at the cellular level, inducing the formation of enormous blebs, particularly after incubation for more than 15 minutes, which were also observed in cells that do not express the LPA_3_ receptor; furthermore, paroxetine also caused the abrupt halting of cyclosis. These data indicate that these events are receptor-independent side effects. However, the expelling of vesicles was only observable in cells that express LPA_3_ receptors. Therefore, the antidepressants examined seem to trigger a complex mixture of receptor-dependent and - independent events. Based on published available data, we can only indicate possible reasons for such effects (next paragraph). Nevertheless, these findings indicate the need to observe what morphological changes occur in the cells and avoid limiting the experiments to the usual pharmacological, biochemical, and molecular biological approaches. These observations open investigation possibilities in cell biology, far from our expertise and technical capacities; hopefully, these effects might interest research groups with such training.

Blebbing is traditionally associated with cell death, but it is now accepted to be also involved in many cellular physiology and pathological events such as division, apoptosis, migration, and many other processes (reviewed in (Charras et al., 2008; Ikenouchi and Aoki, 2022)). This plethora of actions also includes signaling (see, for example (Godin and Ferguson, 2010; Vanderboor et al., 2020)). One of these exciting reviews states: "Blebbing is to a large extent a physical, rather than chemical process, and it occurs over time- and length-scales that have been little investigated in animal cells, especially in the molecular era. It is thus an open and exciting area for biophysical investigation of the forces and mechanics that shape animal cells" (Charras et al., 2008); to the extent of our knowledge, this remains so. The other exciting review indicates: "When the plasma membrane detaches from the underlying actin cortex, the plasma membrane expands according to intracellular pressure and a spherical membrane protrusion, called a bleb, is formed" (Ikenouchi and Aoki, 2022), emphasizing the roles of the cell cytoskeleton in bleb formation; other sections of the same review highlight the function of intracellular calcium and the flexibility of the plasma membrane (Ikenouchi and Aoki, 2022). In this regard, it should be mentioned that imipramine is an inhibitor of Facsin I (Alburquerque-Gonzalez et al., 2020), which is a key protein in actin bundling, playing a causative role in tumor invasion and is overexpressed in different cancer types (Alburquerque-Gonzalez et al., 2020; Li et al., 2022). Therefore, the perturbation of the cytoskeleton dynamics might be involved in the impressive blebbing observed in response to these tricyclic drugs. Antidepressants are, as a group, a highly heterogeneous family of compounds and very promiscuous, i.e., "Velcro-like", meaning that they can interact with and modify the function of many different proteins. At the same time, they are essential therapeutic tools for the treatment of depression and other related psychiatric maladies.

Amitriptyline and paroxetine are known to interact with distinct monoamine reuptake transporters and a variety of receptors, including monoamine receptors such as adrenergic, muscarinic, and those for histamine and serotonin (reviewed in (Bourin et al., 2001; Lawson, 2017)); while imipramine and other antidepressants are known to interact with α_1_-adrenergic receptor subtypes (Chmielarz et al., 2021). The list of antidepressant-interrelating receptors includes those for LPA ((Banks et al., 2018; Chmielarz et al., 2021; Kajitani et al., 2016; Kajitani et al., 2024; Olianas et al., 2015, 2017, 2019, 2020, 2023) and present work). Paroxetine has another important side-effect: the ability to inhibit G protein-coupled receptor kinase 2 (GRK2), which is an essential regulator of G protein-coupled receptor activity and internalization (Homan et al., 2014; Kowalska et al., 2021; Schumacher et al., 2015; Thal et al., 2012). The extent to which these actions participate in the events described here is currently unknown.

It is worth mentioning that some of the side effects of these antidepressants have potential and even clinically tested therapeutic niches; a few of them are indicated next. Paroxetine, through its GRK2 inhibitory activity, triggers several outcomes: renoprotective actions, including histologically determined podocyte protection and reduction of proteinuria in mouse models of diabetic kidney disease (Li et al., 2024); it reverses cardiac dysfunction and remodeling after myocardial infarction in mice (Schumacher et al., 2015); also, in rodents, paroxetine significantly attenuated gland fibrosis and alleviated the progression of Sjögren’s syndrome (Fang et al., 2024); it alters the ability of the active metabolite of fingolimod (i.e., phosphorylated FTY or pFTY) to induce sphingosine 1-phosphate receptor 1 (S1P1) phosphorylation and internalization (Martínez-Morales et al., 2018), suggesting a possible interference with the action of this agent used in multiple sclerosis treatment (Brinkmann et al., 2010). Paroxetine acts directly on beta cells to enhance glucose-stimulated insulin secretion and induces beta-cell mass expansion (Toczyska et al., 2024). Imipramine blocks Non-Small Cell Lung Cancer progression in cultured cells and tumor-bearing animal models (Yueh et al., 2021), and the antitumoral effects of distinct tricyclic drugs in vitro and preclinical studies have been reviewed (Asensi-Canto et al., 2022).

Finally, our present findings shed light on the mechanism of action of antidepressants on the LPA_3_ receptors which likely might participate in their effects on other LPA receptors. It provides clues to decipher the structural changes associated with the different outcomes.

We hope our findings will stimulate the testing of many other agents through docking and experimental studies. Synthesis of compounds with selectivity and defined pharmacodynamic properties is greatly needed for knowledge advance and has therapeutic potential for treating diseases such as ovarian cancer.

## CRediT authorship contribution statement

Conceptualization: K.H.S., J. C.-B., J.A.G.-S.; Methodology: K.H.S., M.T.R.-A., R.R.-H., J.C.-B.;Investigation: K.H.S., M.T.R.-A., R.R.-H., J.C.-B.; Visualization: K.H.S., M.T.R.-A., R.R.-H., J.C.-B.; Writing – original draft: K.H.S., J. C.-B., J.A.G.-S.; Writing – review & editing: K.H.S., M.T.R.-A., R.R.-H., J.C.-B., J.A.G.-S.

## DATA AVAILABILITY STATEMENT

The data are available from the corresponding author upon reasonable request.

## ACKNOWLEDGMENTS

We thank Psicopharma SA de CV (Mexico) for the generous gift of paroxetine and Dr Edith Zárate, Head of Academic Relations and Collaborations of this industry, for her kind help. The advice and technical support of the following members of the indicated Service Units of our Institute is gratefully acknowledged: Dr. Héctor Malagón and Dr. Claudia Rivera (Bioterio); Juan Barbosa and Gerardo Coello (Cómputo); and Aurey Galván and Manuel Ortínez (Taller). K. Helivier Solís is a student of the Programa de Doctorado en Ciencias Biomédicas (UNAM) of Universidad Nacional Autónoma de México (account 520014983).

## SUPPLEMENTARY MATERIAL

**Supplementary Fig. S1.**
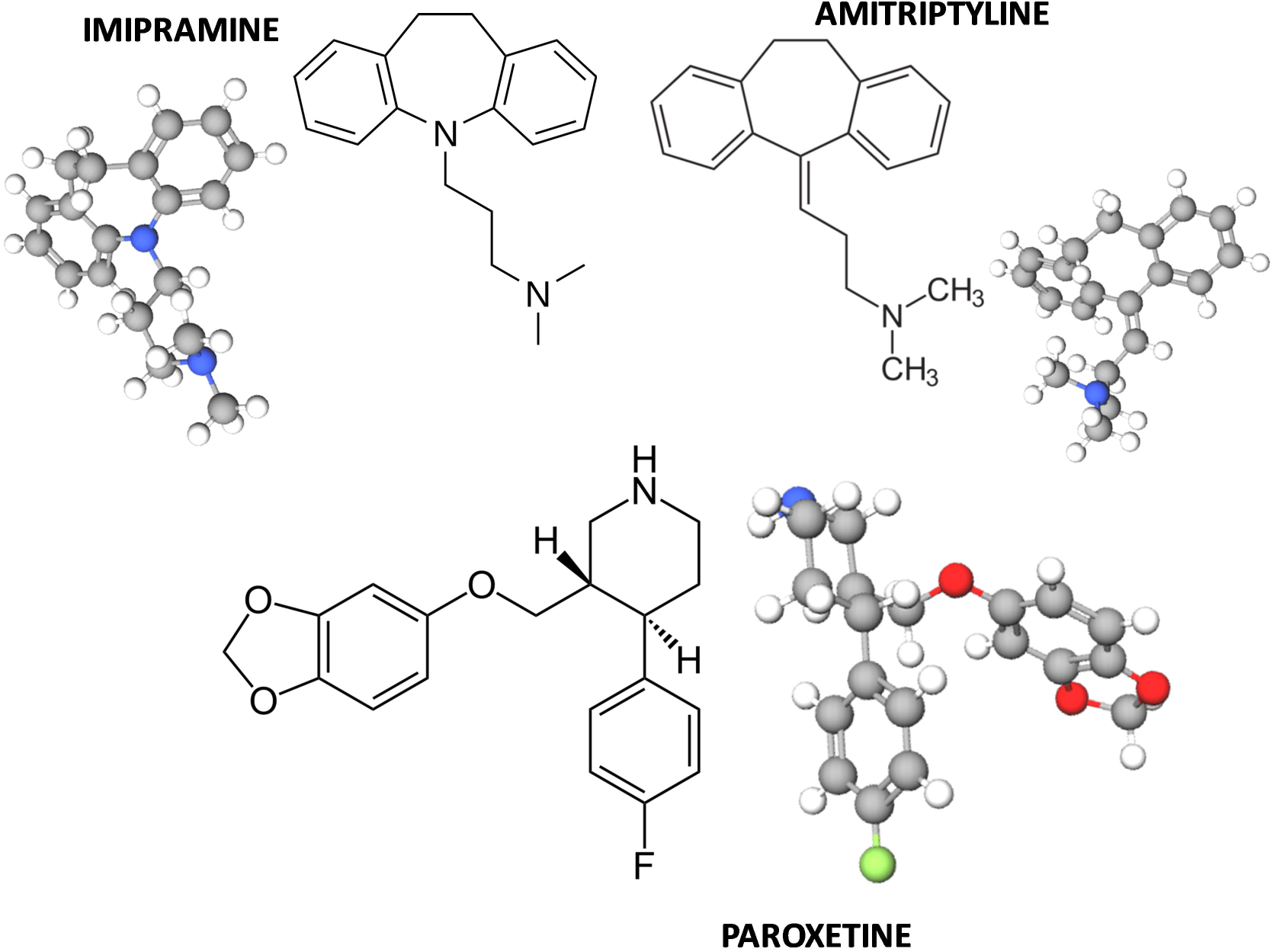
2D and 3D structural formula of the antidepressants imipramine, amitriptyline, and paroxetine. Elements color code: white, hydrogen; gray, carbon; blue, nitrogen; red, oxygen; and green, fluor. Images were obtained using The PyMOL Molecular Graphics System, Version 3.0 Schrödinger, LLC (https://www.pymol.org/support.html?) and BIOVIA; Dassault Systèmes. Discovery Studio Visualizer, v21.1.0.20298; Dassault Systèmes: San Diego, CA, USA, 2021; Available online: https://discover.3ds.com/discovery-studio-visualizer-download (accessed on January 8, 2024).

**Supplementary Fig. S2.**
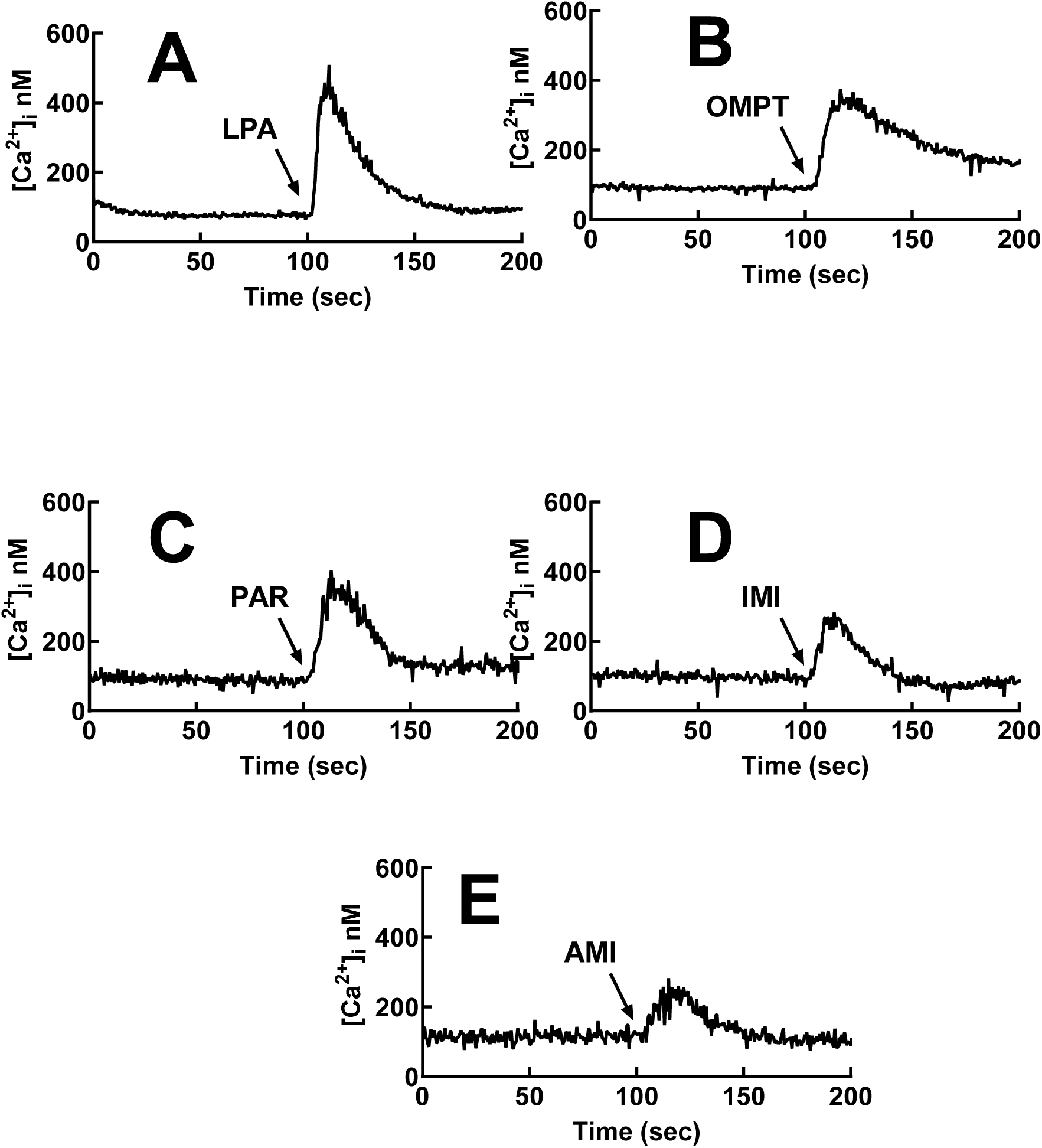
Representative calcium tracings of LPA_3_-expressing cells challenged with 1 µM LPA (A), 1 µM OMPT (B), and 30 µM of the antidepressants: paroxetine (PAR, C), imipramine (IMI, D), and amitriptyline (AMI, E). The arrow indicates the time of addition.

**Supplementary Fig. S3.**
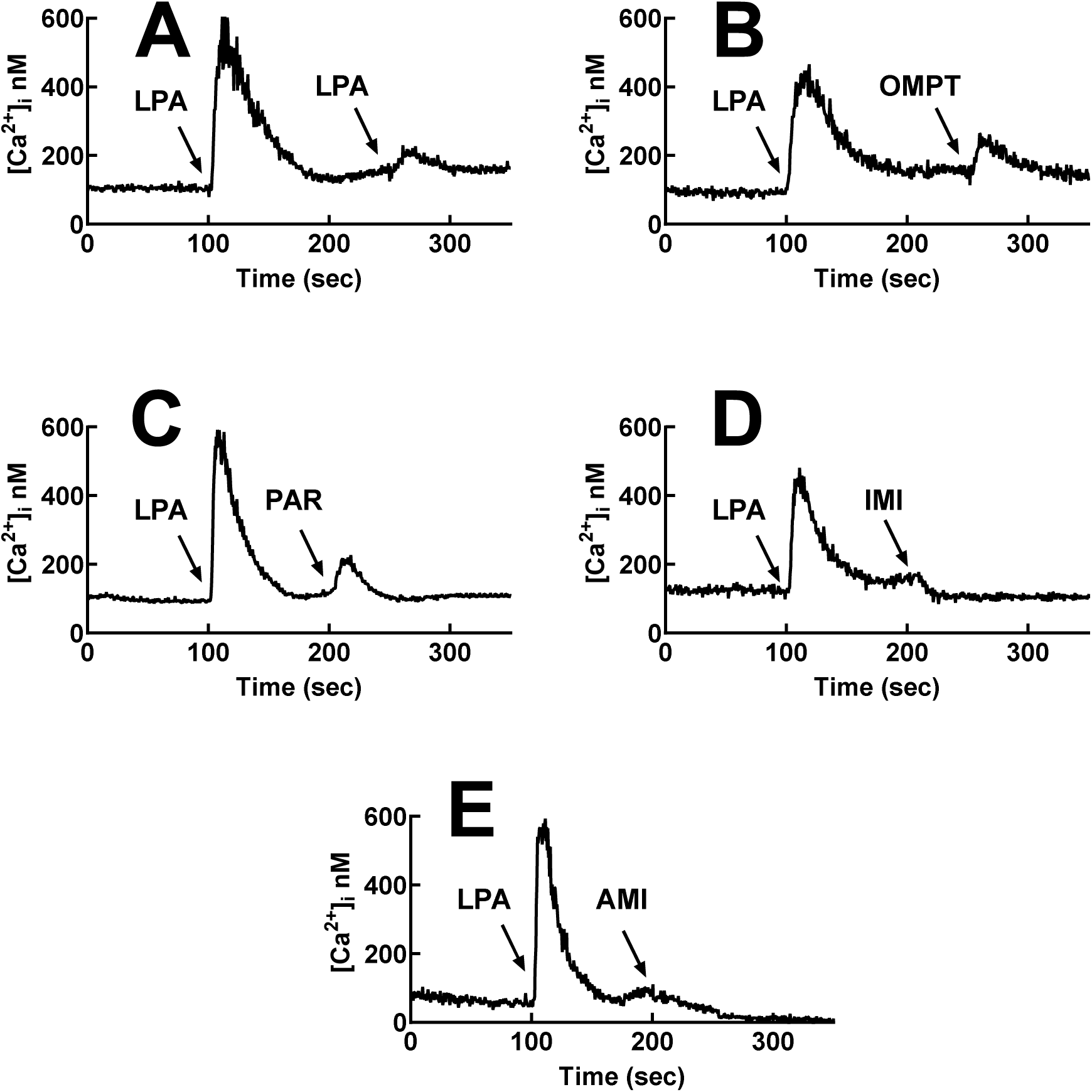
Representative calcium tracings of LPA_3_-expressing cells challenged with 1 µM LPA and then rechallenged with 1 µM LPA (A), 1 µM OMPT (B), and 30 µM of the antidepressants: paroxetine (PAR, C), imipramine (IMI, D), and amitriptyline (AMI, E). Arrows indicate the times of addition.

**Supplementary Fig. S4.**
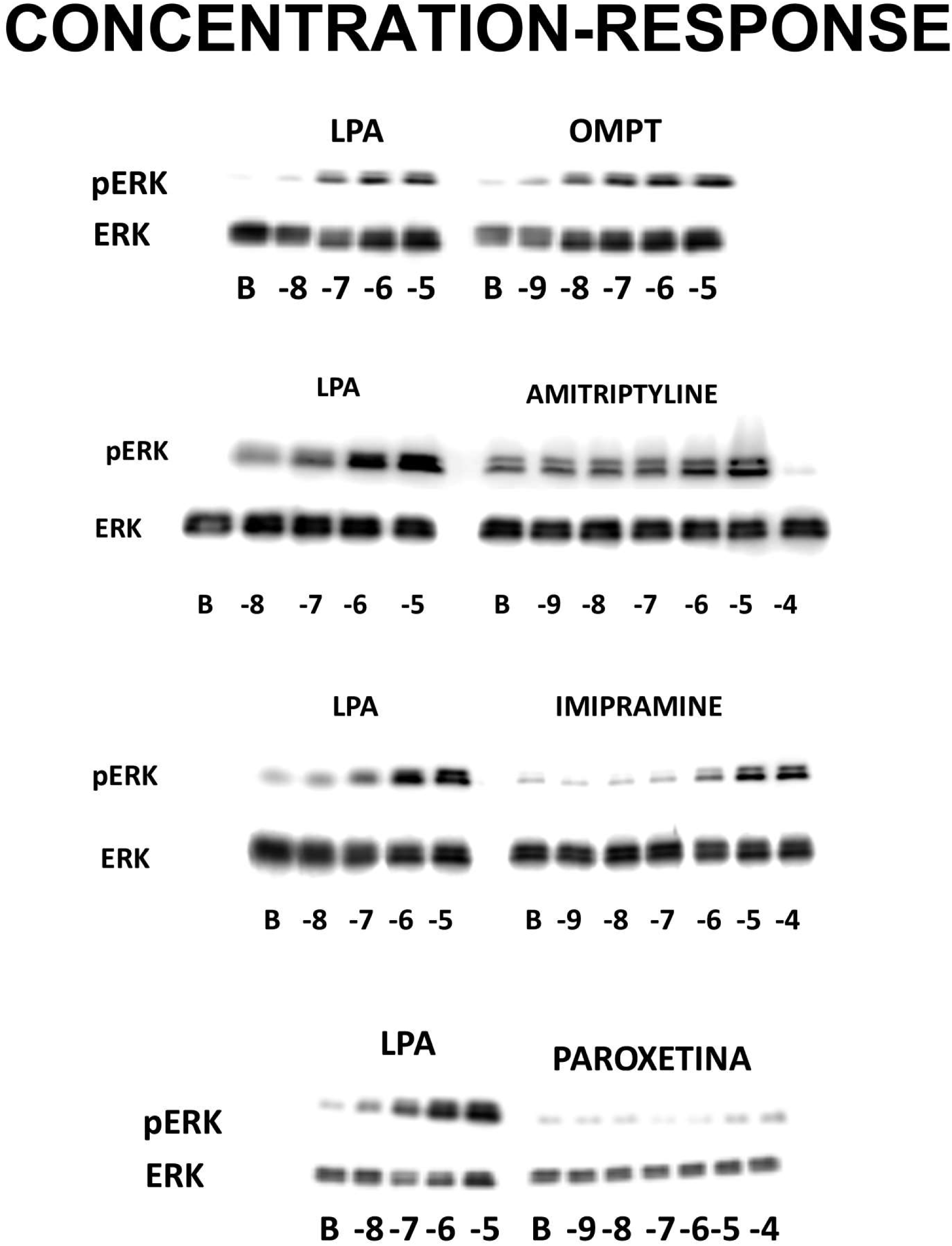
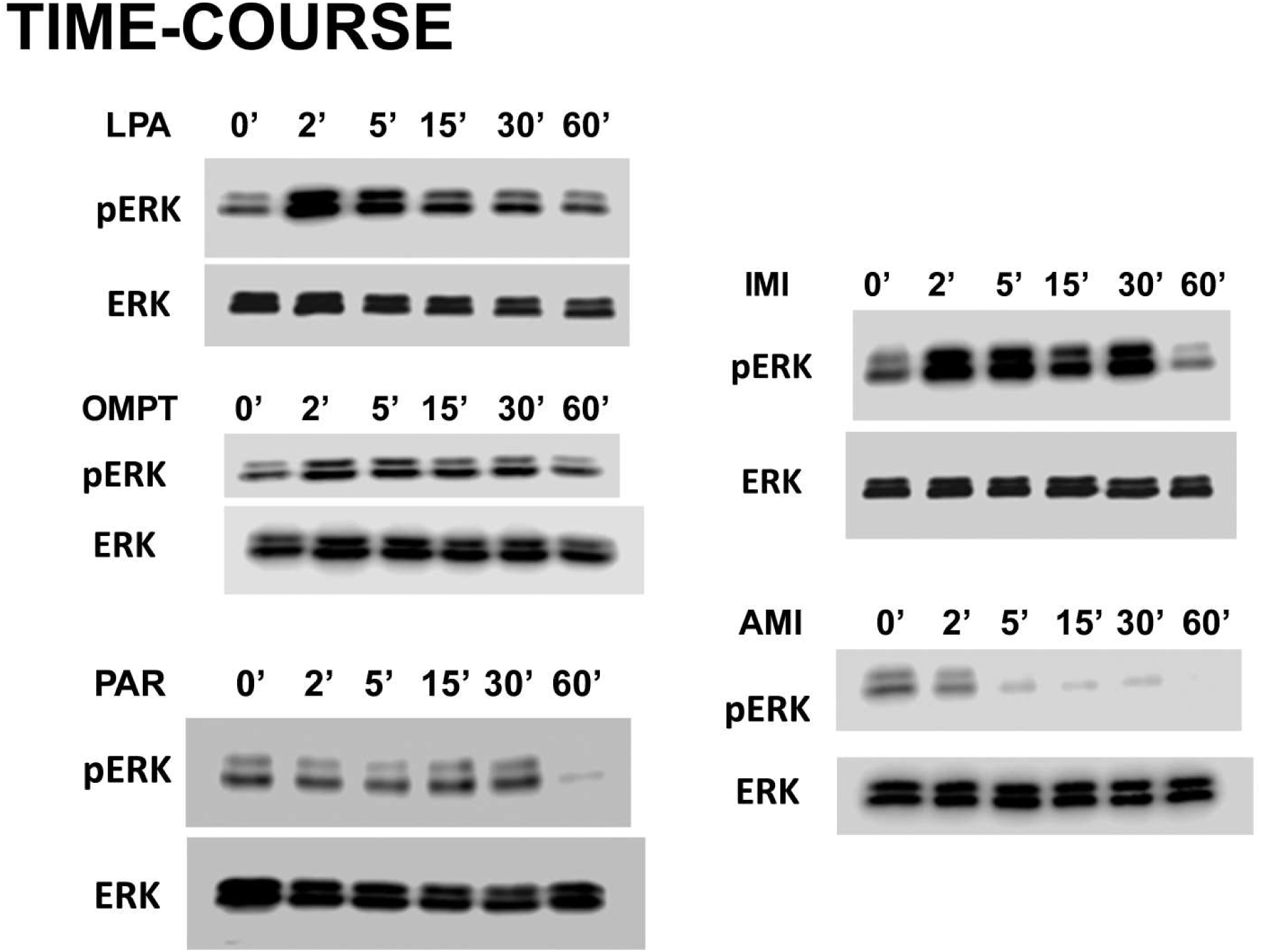
Representative phospho-ERK (pERK) and total ERK Western blots of the action in LPA_3_-expressing cells challenged with LPA, OMPT, paroxetine (PAR), imipramine (IMI), and amitriptyline (AMI). In the upper section (CONCENTRATION-RESPONSE), images corresponding to the concentration of the stimulus (indicated below the blots, M) and the incubation time was 2 min. In the lower section (TIME-COURSE), the time of incubation (min, ’) is indicated above the blots, and the concentration of agents was as follows: 1 µM LPA, 1 µM OMPT, and 30 µM of the antidepressants: paroxetine (PAR), imipramine (IMI), and amitriptyline (AMI).

**Supplementary Fig. S5.**
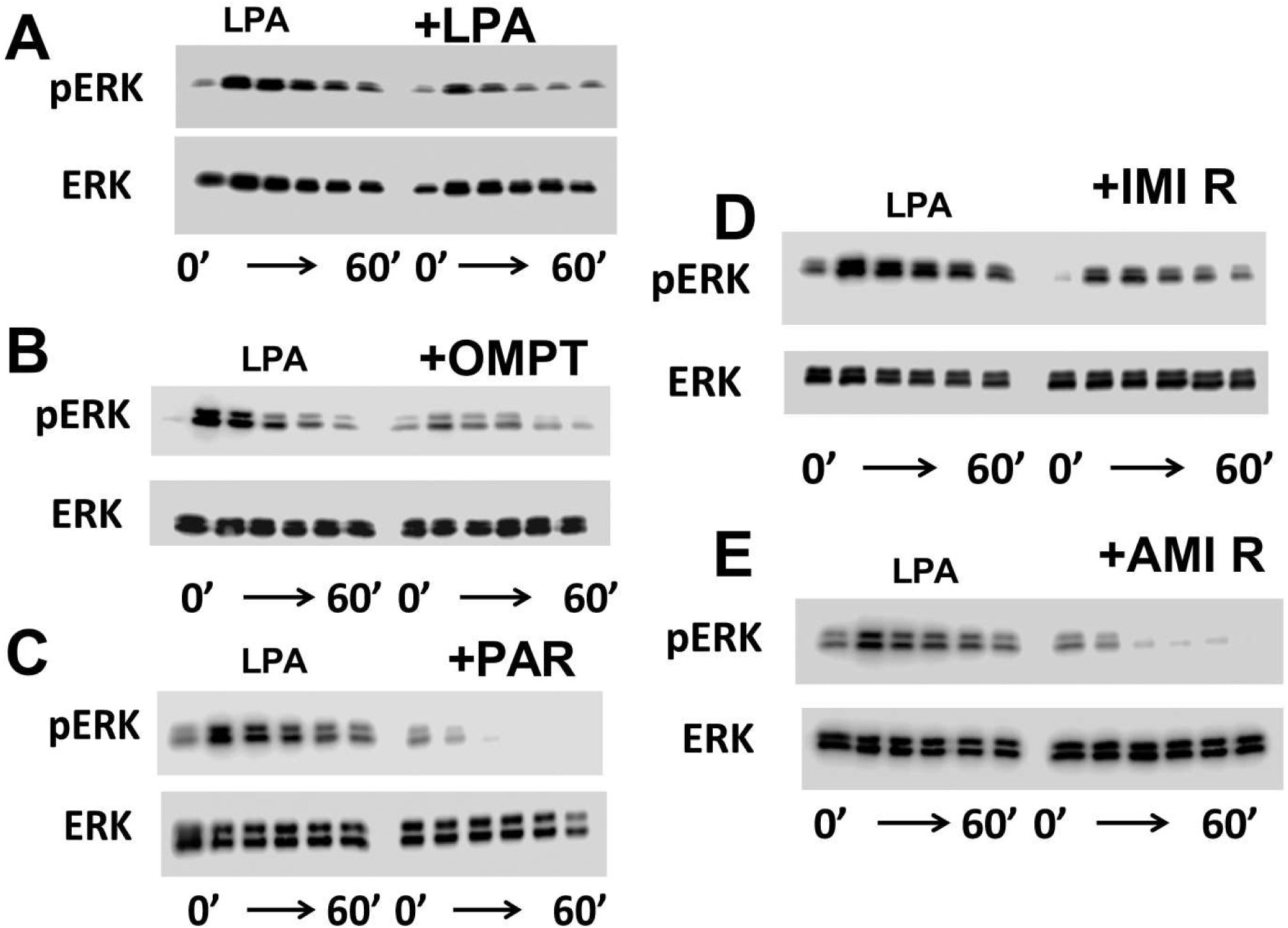
Representative phospho-ERK (pERK) and total ERK Western blots obtained from cells challenged with 1 µM LPA and then rechallenged with 1 µM LPA (A), 1 µM OMPT (B) and 30 µM of the antidepressants: paroxetine (PAR, C), imipramine (IMI, D), and amitriptyline (AMI, E). The time of incubation (min, ’) is indicated below the blots (corresponding to 0, 2, 5, 15, 30 and 60 min). Due to cell detachment (see the manuscript and Supplementary Fig. S6), samples treated with imipramine and amitriptyline contain cells recovered from the incubation medium (indicated as "R").

**Supplementary Fig. S6.**
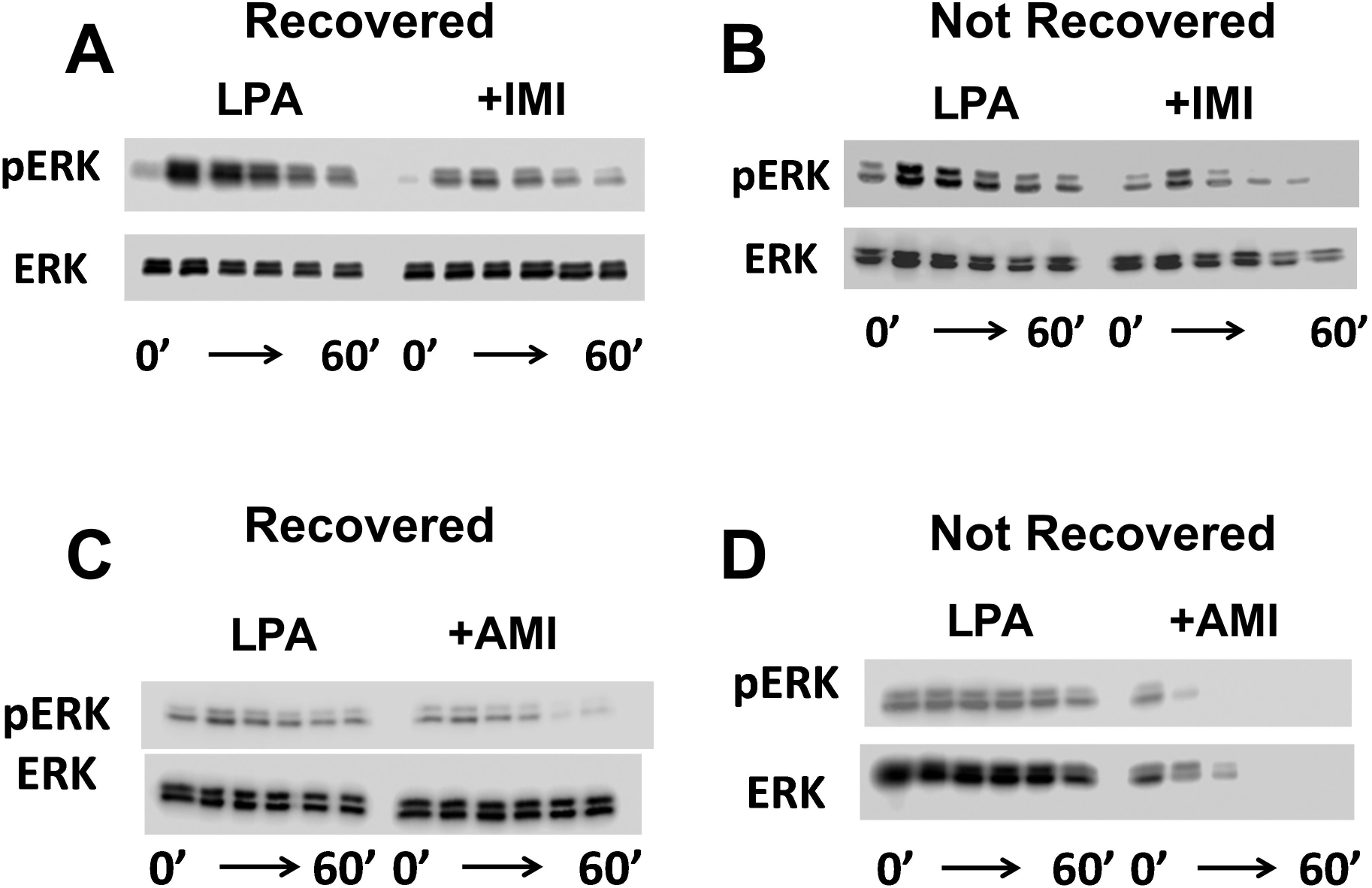
Representative phospho-ERK (pERK) and total ERK Western blots of the action in LPA_3_-expressing cells of cells challenged with 1 µM LPA and then rechallenged with 30 µM, imipramine (IMI, A and B), and amitriptyline (AMI, C and D). The incubation time (min) is indicated below the blots (corresponding to 0, 2, 5, 15, 30 and 60 min). Due to cell detachment (see manuscript text), samples treated with imipramine and amitriptyline contain the cells Recovered (A and C) from the incubation medium or only those remaining attached to the plates (Not Recovered). Notice the material loss in the Not Recovered total ERK samples, particularly those with longer incubation times.

**Supplementary Fig S7.**
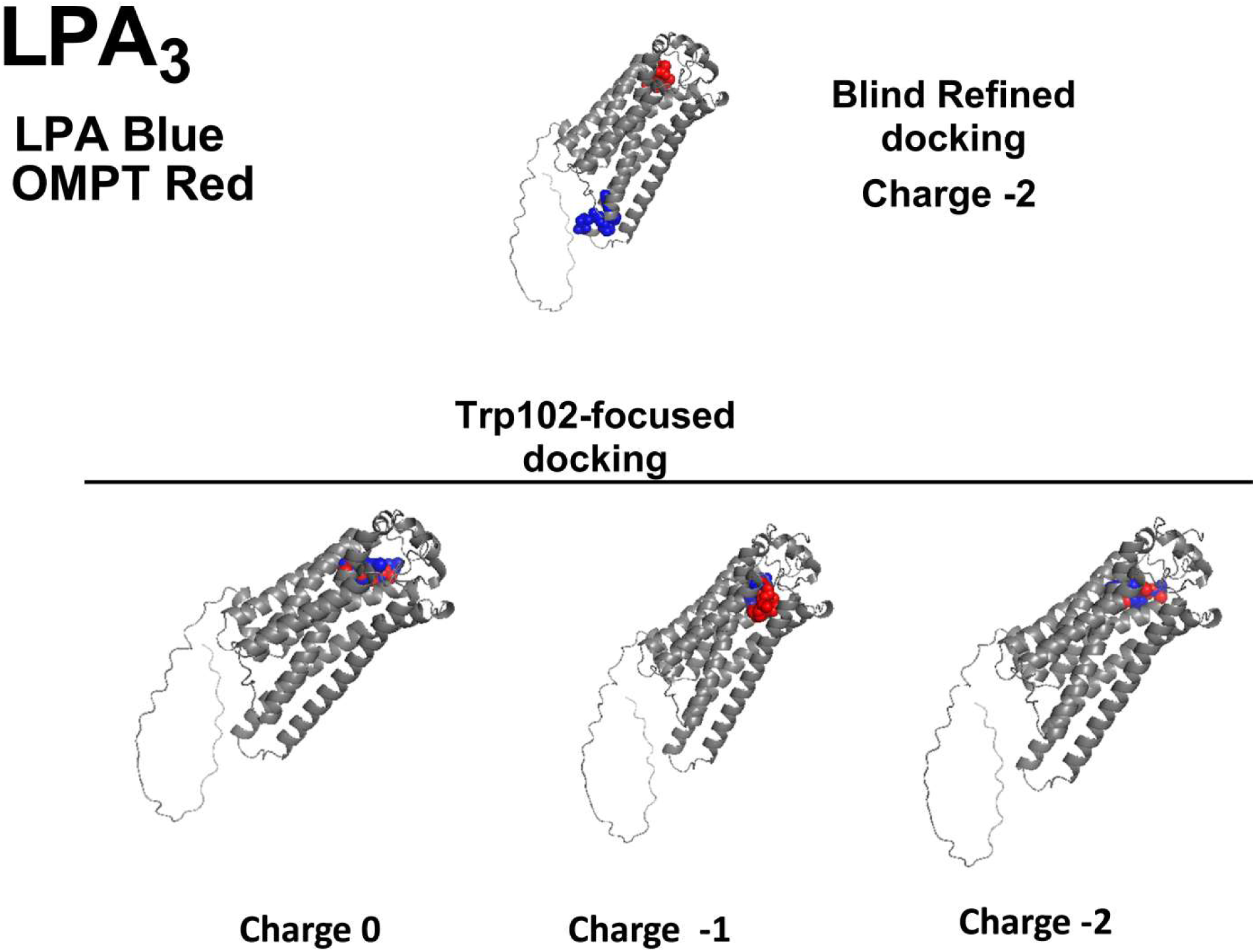
3D Graphical representation of the docking of LPA_3_ receptor to LPA (blue) and OMPT (red). In the figure’s upper part, the Blind refined model was used, and a charge of -2 was used for the ligands. In the figure’s lower part, the Trp102-focused model was employed, and distinct ligand charges are presented. These data were obtained from Solís, K.H., Romero-Ávila, M.T., Rincón-Heredia, R., Correa-Basurto, J., García-Sáinz, J.A., 2025. Multiple LPA3 receptor agonist binding sites evidenced under docking and functional studies. BioRxiv. https://doi.org/10.1101/2024.09.03.611107.

**Supplementary Fig. S8.**
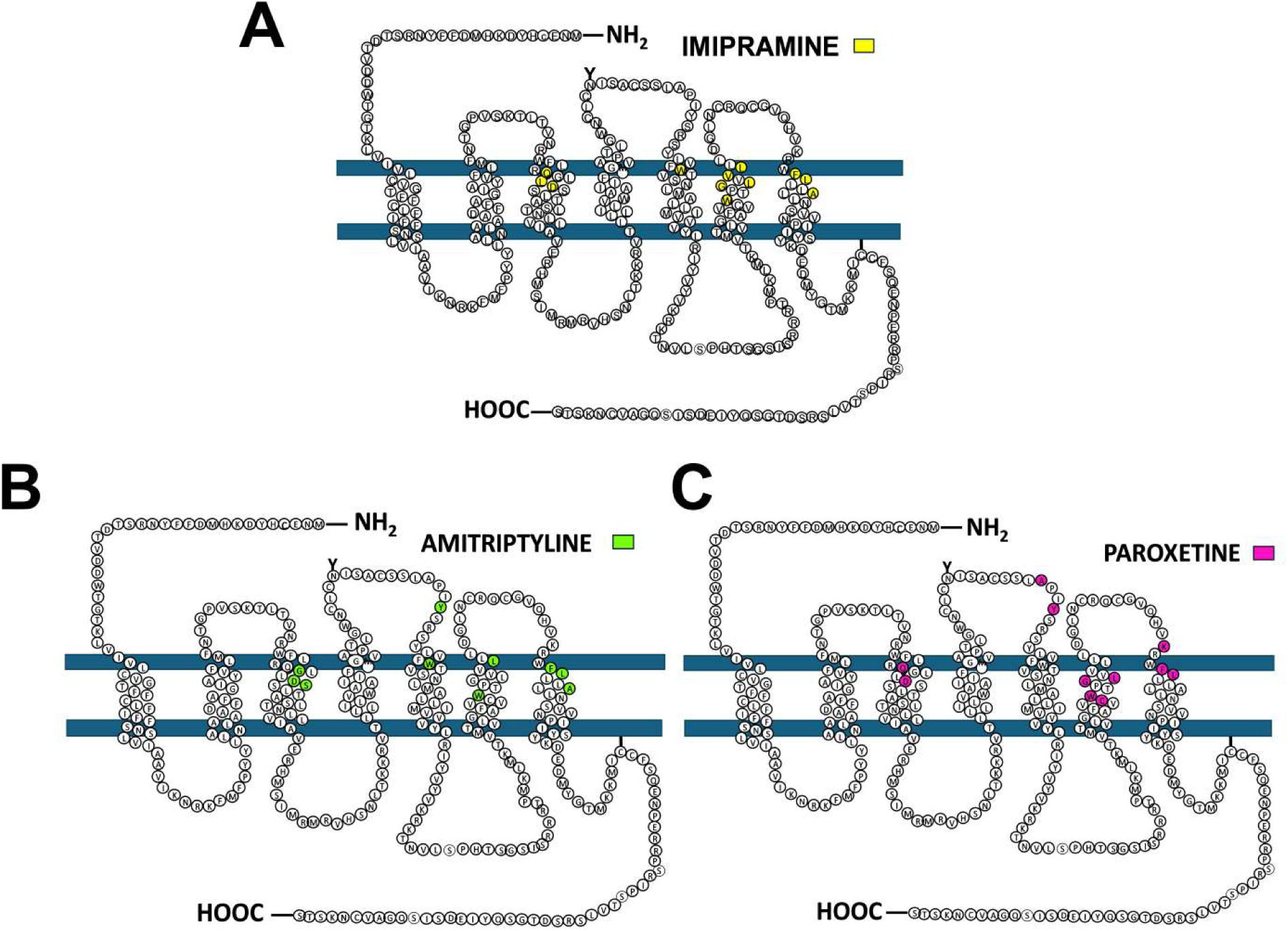
Cartoon representation of the LPA_3_ receptor. The amino acids that interact with imipramine (panel A, yellow), amitriptyline (panel B, green), and paroxetine (panel C, purple) are indicated.

**Supplementary Fig. S9.**
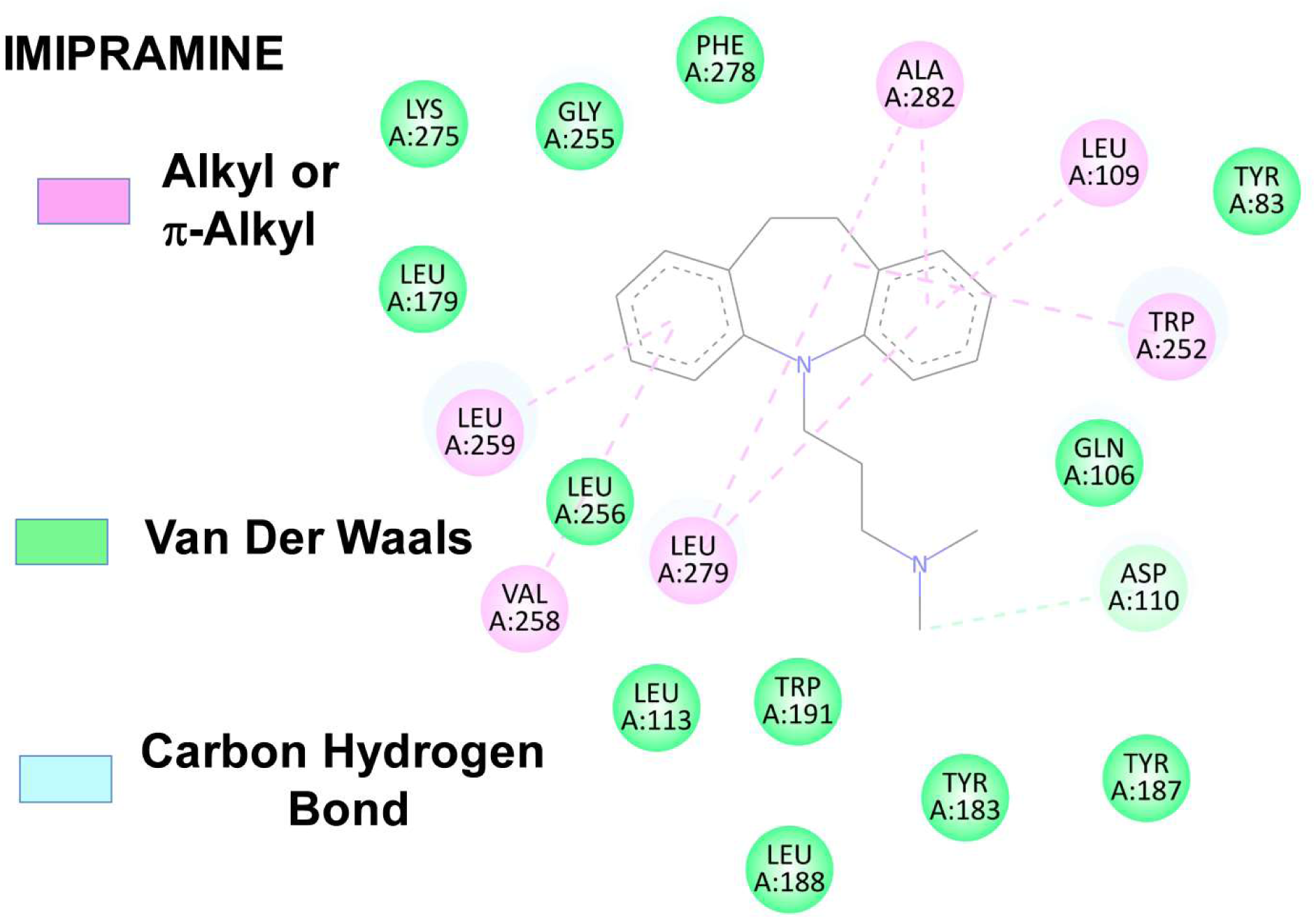
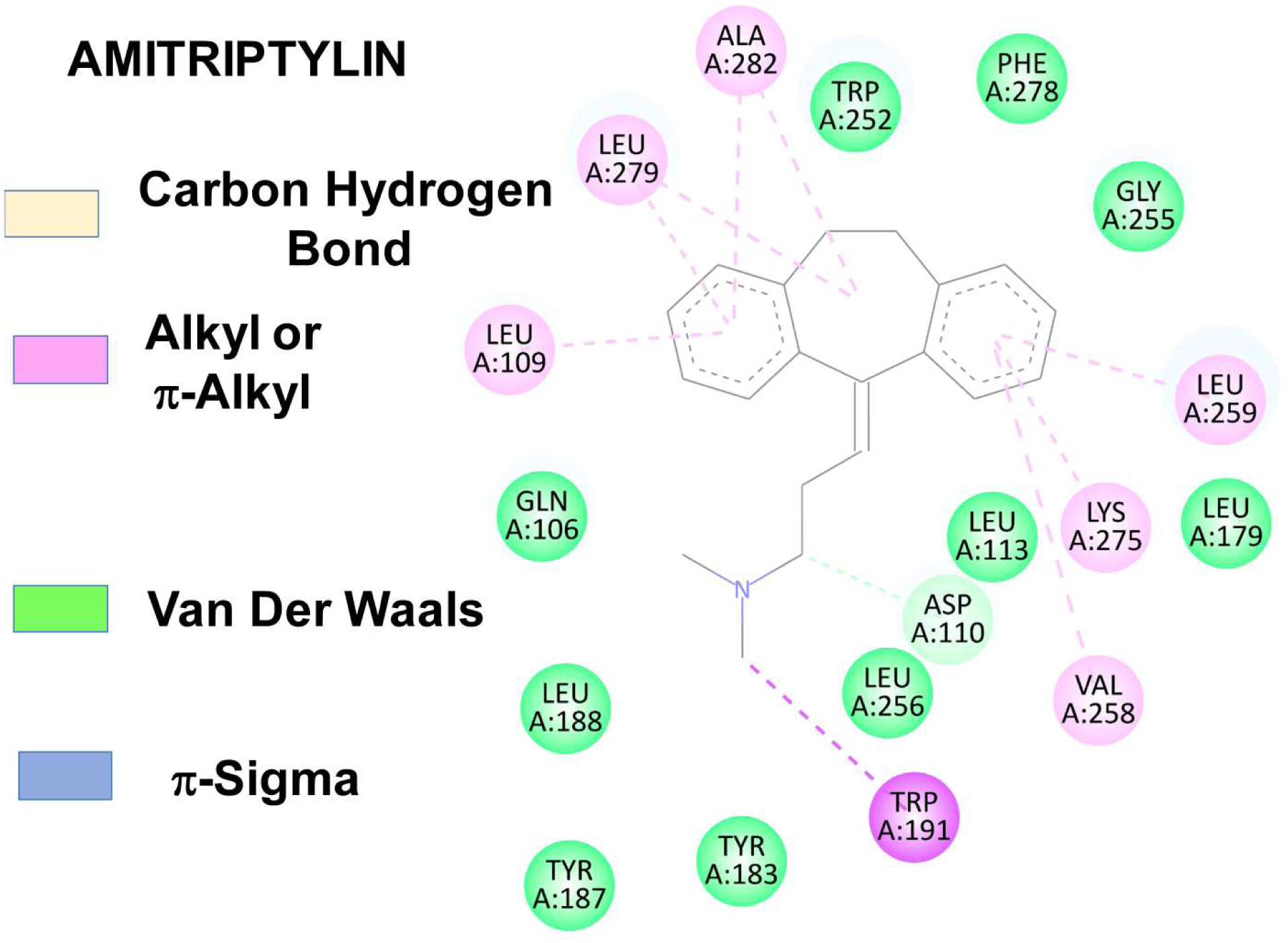
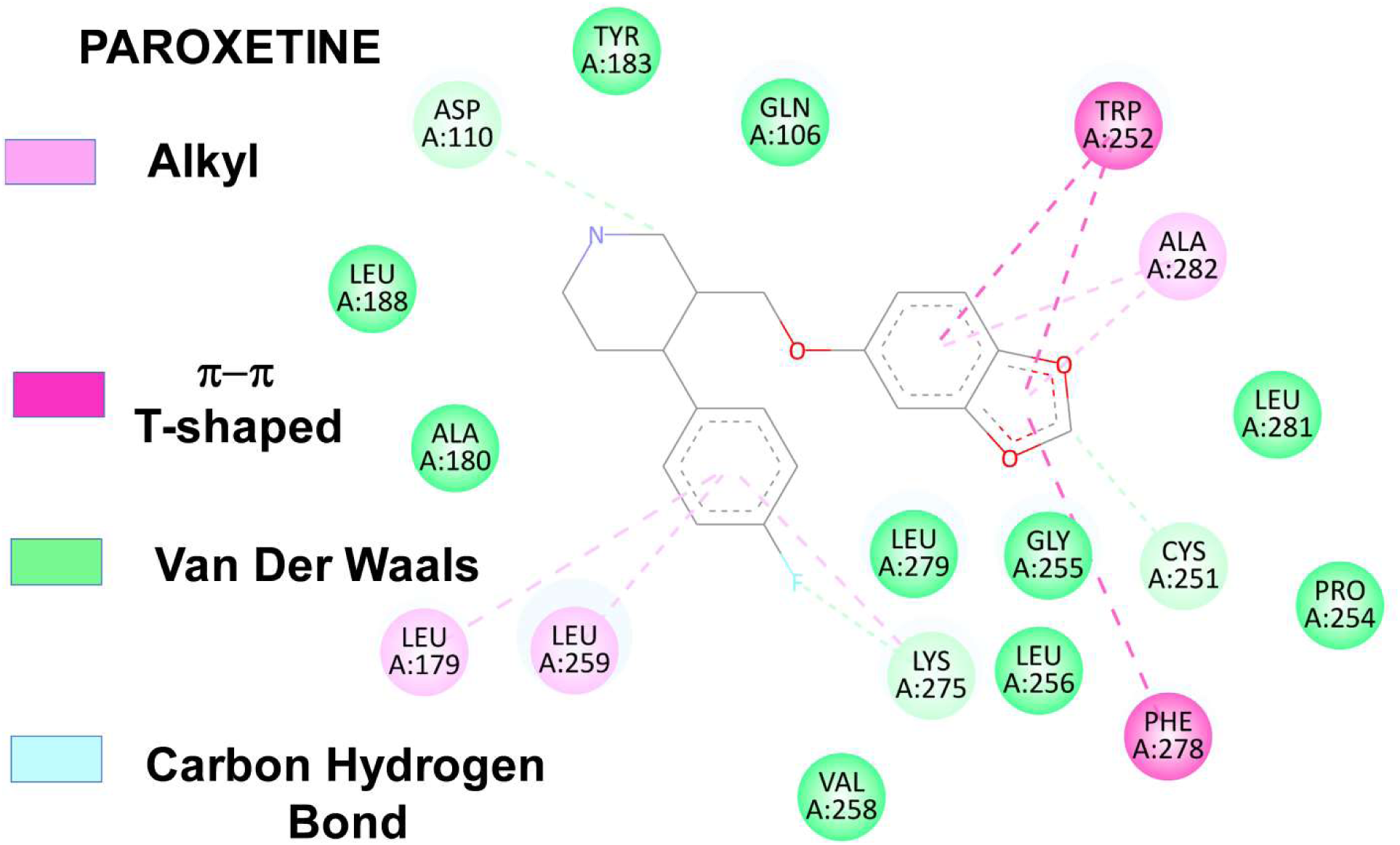
The 2D diagrams, obtained with Discovery Studio Visualizer, show interactions at the LPA_3_ binding site with imipramine (first panel), amitriptyline (second panel), and paroxetine (third panel). Critical amino acids contributing to interactions are shown in circles.

**Supplementary Fig. S10.**
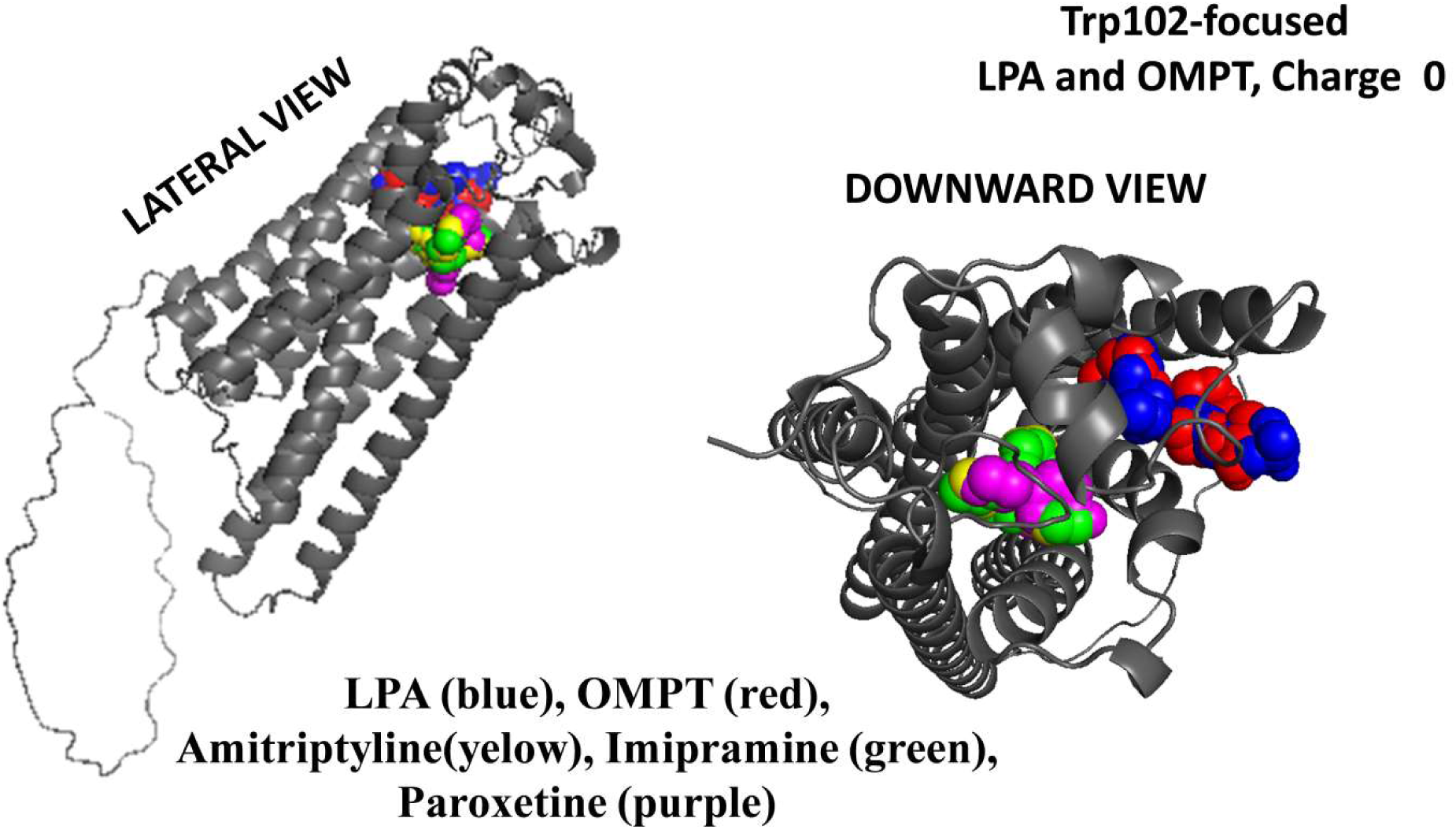
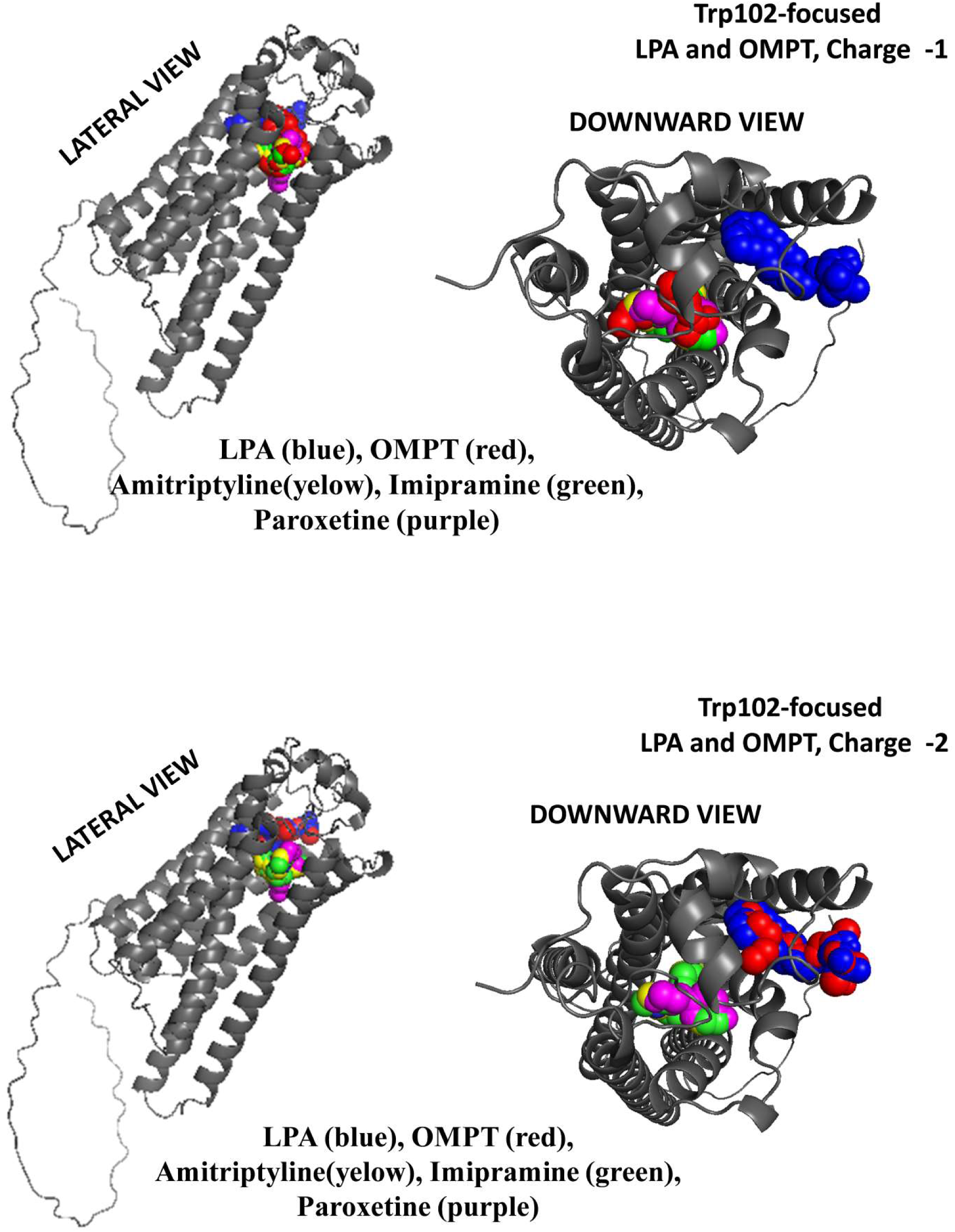
3D Images of the LPA_3_ receptor docking with distinct ligands obtained using PyMol. For LPA and OMPT, the Trp102-focused model was used and the ligand charge used is indicated in the panels. Ligands are color-coded, and both lateral and downward views are included

## VIDEO LEGENDS

**Video 1A Fluorescence. Representative video of paroxetine action on LPA_3_ expressing cells.** Fluorescence (receptors) delineated the plasma membrane and it was also located in vesicles inside the cells. Almost immediately after paroxetine addition, fluoresce coalesce forming patches that now indicate the location of the plasma membrane in a “dotted-line fashion”. It can also be observed that cyclosis abruptly diminished near to a full-stop, although some changes in cell shape were noticed.

**Video 1B Merged Fluorescence/ DIC (differential interference contrast). Representative video of paroxetine action on LPA_3_ expressing cells.** This merged-images video allows observation of the marked changes in the shape and cell surface dynamics. It can also be more clearly observed that cyclosis abruptly stopped but that movement of cell debris continued. In addition “expelling” of dense vesicles that remain in proximity to the cells can also be appreciated.

**Video 2A Fluorescence. Representative video of amitriptyline action on LPA_3_ expressing cells.** After a short delay amitriptyline addition induced some cell contraction and the formation of an enormous amount of blebs of large size. Fluorescence accumulates inside the cells in discrete cell regions and the cells reorganize appearing to be inside gigantic blebs. At a given point the two central cells seem to “fuse”, appearing to be both inside a single cell blister.

**Video 2B Merged Fluorescence/ DIC. Representative video of amitriptyline action on LPA_3_ expressing cells.** This merged-images video allows observation of the changes described for Video 2B but in addition appears to show that a “space” with few organelles exist between the enormous bleb that surrounds each cell and what seems to be compacted cells. In addition the apparent fusion of the cells is more clearly observed. Appearance of vesicles with optically dense material inside and outside the cells was also detected at longer times of incubation.

**Video 3A Fluorescence. Representative video of imipramine action on LPA_3_ expressing cells.** Imipramine (as amitriptyline addition did) induced cell contraction and the formation of blebs of large size. The actions of this tricyclic antidepressant also lead cell organelles to condensate appearing to be inside gigantic blebs.

**Video 3B Merged Fluorescence/ DIC. Representative video of imipramine action on LPA_3_ expressing cells.** This video allows a more clear observation of the bleb formation and fusion to be gigantic and apparently “engulf” most of the cell whose organelles seem to concentrate and then “swell” (i.e., increase its apparent volume). Vesicles expelling was not evidenced with this agent.

**Video 4 DIC. Representative video of paroxetine action on uninduced cells.** This video evidenced that paroxetine induced rapid cell contraction, bleb formation and an abrupt interruption of cyclosis in the absence of LPA_3_ receptor expression (i.e., these action are receptor-independent). In contrast, we did not observe any vesicle expel, which suggests that this event is receptor-mediated.

**Video 5 DIC. Representative video of amitriptyline action on uninduced cells.** This video evidenced that amitriptyline induced rapid cell contraction, and marked bleb formation in the absence of LPA_3_ receptor expression (i.e., receptor-independent actions).

**Video 6 DIC. Representative video of imipramine action on uninduced cells.** This video evidenced that imipramine induced rapid cell contraction, and marked bleb formation in the absence of LPA_3_ receptor expression (i.e., receptor-independent actions).

